# Multi-omic profiling reveals widespread dysregulation of innate immunity and hematopoiesis in COVID-19

**DOI:** 10.1101/2020.12.18.423363

**Authors:** Aaron J. Wilk, Madeline J. Lee, Bei Wei, Benjamin Parks, Ruoxi Pi, Giovanny J. Martínez-Colón, Thanmayi Ranganath, Nancy Q. Zhao, Shalina Taylor, Winston Becker, Stanford COVID-19 Biobank, David Jimenez-Morales, Andra L. Blomkalns, Ruth O’Hara, Euan A. Ashley, Kari C. Nadeau, Samuel Yang, Susan Holmes, Marlene Rabinovitch, Angela J. Rogers, William J. Greenleaf, Catherine A. Blish

**Author notes:** These authors contributed equally to this work (Aaron J. Wilk, Madeline J. Lee, Bei Wei, Benjamin Parks). Correspondence to: William J. Greenleaf, Catherine A. Blish.

## Abstract

Our understanding of protective vs. pathologic immune responses to SARS-CoV-2, the virus that causes Coronavirus disease 2019 (COVID-19), is limited by inadequate profiling of patients at the extremes of the disease severity spectrum. Here, we performed multi-omic single-cell immune profiling of 64 COVID-19 patients across the full range of disease severity, from outpatients with mild disease to fatal cases. Our transcriptomic, epigenomic, and proteomic analyses reveal widespread dysfunction of peripheral innate immunity in severe and fatal COVID-19, with the most profound disturbances including a prominent neutrophil hyperactivation signature and monocytes with anti-inflammatory features. We further demonstrate that emergency myelopoiesis is a prominent feature of fatal COVID-19. Collectively, our results reveal disease severity-associated immune phenotypes in COVID-19 and identify pathogenesis-associated pathways that are potential targets for therapeutic intervention.

**One Sentence Summary:** Single-cell profiling demonstrates multifarious dysregulation of innate immune phenotype associated with COVID-19 severity.

## INTRODUCTION

The COVID-19 pandemic, caused by the severe acute respiratory syndrome coronavirus 2 (SARS-CoV-2), is an urgent public health crisis. COVID-19 has a highly variable disease course: approximately 20% of infected individuals require hospitalization, ~5% require critical care, and as many as 40% of cases are asymptomatic *(1*, *2)*. As the immune response to SARS-CoV-2 is a key determinant of COVID-19 severity and outcome, understanding the immunological underpinnings of COVID-19 pathogenesis is critical to predict, prevent, and treat SARS-CoV-2 infection and to prepare for the possibility of future infections caused by emerging betacoronaviruses that may be introduced from existing reservoirs *(3*–*6)*.

Severe COVID-19 is associated with a number of distinct immunological signatures. For example, increased serum levels of proinflammatory cytokines such as IL-1β, IL-6, IP-10, and TNFα and the alarmins S100A8 and S100A9 are associated with worse outcomes *(7*–*12)*. Patients with less severe disease tend to have both lower levels of proinflammatory cytokines and higher levels of tissue repair factors *(9)*. COVID-19 also reconfigures leukocyte phenotype in a severity-specific fashion, with severe COVID-19 associated with lymphocyte exhaustion *(13*–*15)*, neutrophil activation signatures *(16*–*20)*, and hematopoietic alterations *(15*, *21)*. While many of these findings have been established through transcriptomic and proteomic profiling, the gene regulatory changes underlying severe disease manifestations have not been determined.

Comparatively less is known about the features of immune responses to SARS-CoV-2 that protect against severe disease, as most cohorts profiled to date have included only hospitalized patients. Neutralizing antibodies and virus-specific T cell responses have been detected in mildly symptomatic patients, providing evidence of a successful adaptive immune response across the disease spectrum *(22*–*27)*. Notably, patients with mild COVID-19 have much lower levels of proinflammatory plasma cytokines, suggesting that the immune response in mild disease can eradicate the virus without triggering the hyperinflammatory state observed in severe cases *(9*, *11)*. Therefore, to define protective versus pathological features of the immune response, we aimed to profile both mild (WHO score 1-3, no oxygen requirement), moderate (WHO score 4-5, noninvasive oxygen requirement), and severe (WHO score 6-8, intubated) cases of COVID-19.

To map the immune response at the epigenetic, transcriptional, and proteomic level, we performed single-cell assay for transposase-accessible chromatin (scATAC-seq), single-cell RNA-sequencing (scRNA-seq), and cytometry by time-of-flight (CyTOF) on the peripheral immune cells of a cohort of COVID-19 patients across the entire spectrum of disease severity. We discovered many immunological perturbations associated with disease severity, including robust signatures related to neutrophil activation along with dysfunction of monocytes, type 2 conventional dendritic cells, and natural killer (NK) cells. In addition, we found strong evidence for emergency myelopoiesis in fatal disease. We also identified epigenetic changes correlated to these transcriptional and proteomic changes, demonstrating coordinated changes in regulatory element accessibility and transcription at key pro-inflammatory cytokine-encoding genes in monocytes. Together, this dataset reveals novel mechanistic insights into the pathological and protective mechanisms of the immune response to SARS-CoV-2.

## RESULTS

### A trimodal single-cell atlas of the peripheral immune response to SARS-CoV-2

To investigate how immune responses vary between different severities of COVID-19, we profiled peripheral blood immune cells from 64 patients with COVID-19 and 12 healthy controls with three high-dimensional single-cell modalities: Seq-Well *(28*, *29)*-*based* scRNA-seq (33 patients, 8 controls; including 7 patients previously profiled by our group *(15)*), scATAC-seq (18 patients, 7 controls), and CyTOF (64 patients, 12 controls; **Fig. 1A, Fig. S1A-B**). Importantly, we profiled COVID-19 patients across the full range of the disease severity spectrum, including patients with mild disease (WHO score 1-3, see **Methods**) and hospitalized inpatients with moderate disease (WHO score 4-5) as well as critically ill patients with severe disease (WHO ordinal score 6-8). We scored patients by both peak severity (denoted by the colors of cells/patients) and severity at time of sample collection (separated as groups for box plots). The median age of profiled patients was 43.5 years and 51% were female (**Fig. 1B, Fig. S1B, Table S1**). The majority of patients were sampled during the acute phase of infection; 13 mild and moderately-ill patients were sampled in the convalescent phase (>21 days after first positive nasopharyngeal swab). Demographic information, additional clinical metadata, and the modalities applied to each sample are available in **Table S1**.

**Fig. 1.**
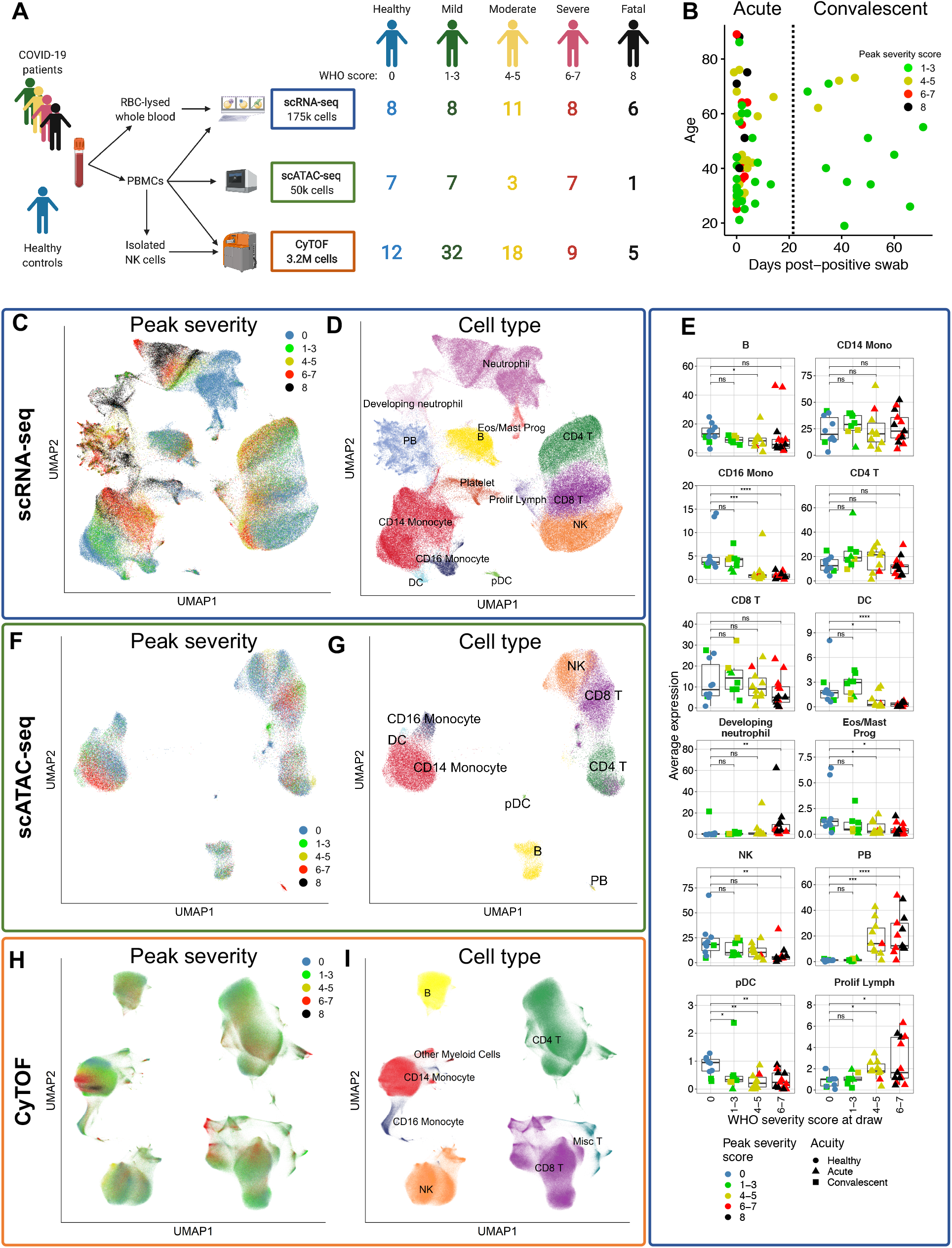
A trimodal single-cell atlas of the peripheral immune response to COVID-19 across a range of disease severities. **A)** Pipeline for sample processing and number of patients analyzed summarized by modality and peak disease severity score. For all display figures, scRNA-seq-derived data are boxed in blue, scATAC-seq-derived data are boxed in green, and CyTOF-derived data are boxed in orange. **B)** Summary of key patient metadata, including age, peak disease severity score, and days after first positive nasopharyngeal (NP) PCR test. The vertical dotted line placed at 21 days post-positive test indicates the threshold after which patient samples are considered convalescent. **C-D, F-I)** UMAP projections of complete scRNA-seq (**C-D**), scATAC-seq (**F-G**), and CyTOF (**H-I**) datasets colored by peak disease severity score of sample (**C,F,H**) or cell type (**D,G,I**). **E)** Cell type proportions from scRNA-seq data of PBMCs in each sample are colored by peak disease severity score. Platelets are excluded from the proportion calculations, as their presence is related to sample processing. The *x* axes correspond to the disease severity score for each sample at the time of collection. *, p < 0.05; **, p < 0.01; ***, p < 0.001; ****, p < 0.0001, n.s., not significant at p = 0.05 by two-sided Wilcoxon rank-sum test with Bonferroni’s correction for multiple hypothesis testing.

Peripheral blood mononuclear cells (PBMCs) were sampled by all modalities; additionally, we processed red blood cell-lysed whole blood by scRNA-seq to profile granulocytes like neutrophils (see **Fig. 5**), and we processed isolated NK cells by CyTOF with a panel enabling deep interrogation of NK cell receptor expression (see **Fig. 4**; **Fig. 1A**). In total, we analyzed ~175,000 single transcriptomes, ~50,000 single chromatin accessibility profiles, and >3.2 million single proteomic profiles (**Fig. 1A, Tables S2-5**). After performing modality-specific quality control procedures (see **Methods**), we created a merged feature matrix of all profiled samples which we subjected to dimensionality reduction, graph-based clustering, and cell type annotation (**Fig. 1C-I, Fig. S1C**, see **Methods, Table S6**).

We first examined how COVID-19 impacted the composition of peripheral immune cells. We saw similar trends in immune cell composition between the three modalities, including depletion of CD16 monocytes, dendritic cells (DCs), and natural killer (NK) cells, as well as increases in plasmablasts in patients with severe and fatal COVID-19 (**Fig. 1E; Fig. S2-3**). Notably, cell subset proportions that were altered in moderate and severe disease were generally unchanged in mild cases, with the exception of plasmacytoid DCs (pDCs), which were depleted in all severity groups (**Fig. 1E; Fig. S2-3**). Further, a population of developing (or immature) neutrophils first identified in our prior study of seven patients *(15)* was confirmed in 17 additional patients (**Fig. 1G, Fig. S1C**) and is similar to that observed by other groups *(21)*.

### Multimodal reference mapping enables accurate annotation and analysis of cellular subtypes

Accurate identification of fine-grained immune cell subtypes is crucial to understanding how COVID-19 reconfigures the immune system; however, these cell types can be difficult to identify *de novo* in single-cell RNA-seq data due to data sparsity and lack of information on canonical surface marker expression in each cell. To address this, we mapped our transcriptomic dataset to a large multimodal reference dataset introduced by Seurat v4, which incorporated extensive surface marker information to improve cell type calls (**Fig. 2A-B, Fig. S4A-B**) *(30)*.

**Fig. 2.**
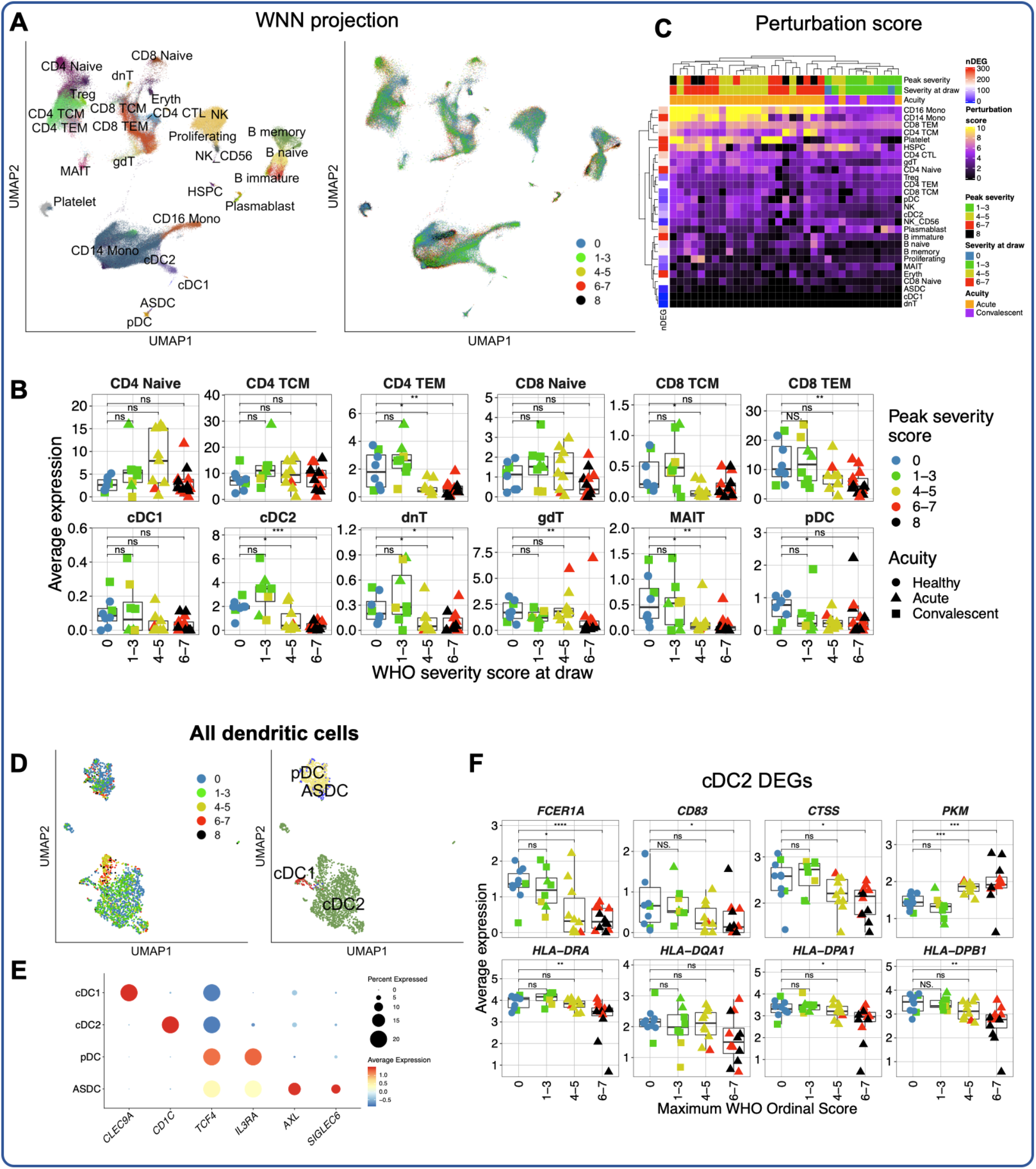
Reference-based cell subtype annotations reveal disease severity-associated perturbations in immune cell subtypes. **A)** Weighted-nearest neighbors projection of scRNA-seq dataset colored by cell type labels transferred from Seurat v4 (left), or by peak disease severity score (right). **B)** Selected cell type proportions in scRNA-seq dataset of cell subtype labels from Seurat v4. Neutrophils and developing neutrophils are excluded from calculation of cell type proportions as these cells do not appear in the Seurat v4 reference. **C)** Heatmap of cellular perturbation scores, as described by Papalexi, et al. *(31)*, per COVID-19 sample in each Seurat v4-labeled cell type. The number of DEGs between all COVID-19 cells and healthy cells for each cell type is plotted at the left. **D)** UMAP projection of all DC subsets colored by peak disease severity score (left) and Seurat v4-annotated cell type (right). **E)** Dot plot depicting percent and average expression of canonical DC genes defining the four annotated DC subsets (see **Methods**, **Fig. S4A**). **F)** Box plots depicting average expression of selected DEGs (see **Table S9** for complete list) by cDC2 cells for each sample. For all box plots: points are colored by the peak disease severity score, shaped according to disease acuity, and grouped by the disease severity score at time of sample collection. *, p < 0.05; **, p < 0.01; ***, p < 0.001; ****, p < 0.0001, n.s., not significant at p = 0.05 by two-sided Wilcoxon rank-sum test with Bonferroni’s correction for multiple hypothesis testing.

Alignment to the Seurat v4 reference dataset revealed gene expression profiles of cell subtypes matching their expected biological signatures (**Fig. S4A-B, Tables S7-8**). For example, multiple T cell subsets, including γδ T cells, mucosal-associated invariant T (MAIT) cells, and regulatory T cells (Tregs) were revealed with Seurat v4 at the expected proportions (**Fig. 2B**) and with the expected transcriptomic phenotype (eg. highly-specific expression of *TRDC* and *TRGC1* by γδ T cells, and *FOXP3* by Tregs; **Fig. S4A**). These annotations closely matched the manually-generated labels; importantly, cell types present in our dataset but absent from the reference (ie. neutrophils) were not successfully mapped (**Fig. S4C,D**). To orthogonally confirm the accuracy of these annotations, we compared the abundances of MAITs measured by CyTOF to the proportions of MAITs in the transcriptomic dataset predicted by Seurat v4. This analysis revealed high concordance between modalities, supporting the accuracy of these annotations (**Fig. S4E**).

This strategy revealed previously unreported changes in peripheral immune cell composition in COVID-19, including the depletion of both CD4 and CD8 T effector memory (T_EM_) cells and double-negative T cells in severe and fatal COVID-19 (**Fig. 2B**). While DCs are known to be depleted in COVID-19, our analysis reveals that this was due to depletion of the cDC2 subset, while cDC1 cells were unchanged (**Fig. 2B**). To prioritize downstream analysis of these cell subsets we calculated a perturbation score *(30*, *31)* for each cell type from each COVID-19 sample relative to healthy controls (see **Methods**). The perturbation score for each cell type is calculated by first identifying genes that display evidence of differential expression between COVID-19 samples and healthy controls, calculating the difference of pseudobulk expression vectors of these genes between COVID-19 samples and healthy controls, and finally projecting the whole transcriptome of each donor onto this vector. This score therefore represents the magnitude of whole transcriptome shifts in gene expression, and reveals disease severity-associated patterns in cell subtype perturbation (**Fig. 2C**). This perturbation score captured phenotypic changes in major cell types like monocytes (explored further in **Fig. 3**) as well as more granular subtypes like CD8 T_EM_ cells. We focused our downstream analysis on cell subtypes with COVID-19 severity-associated perturbation with a high number of DEGs relative to other cell subtypes.

**Fig. 3.**
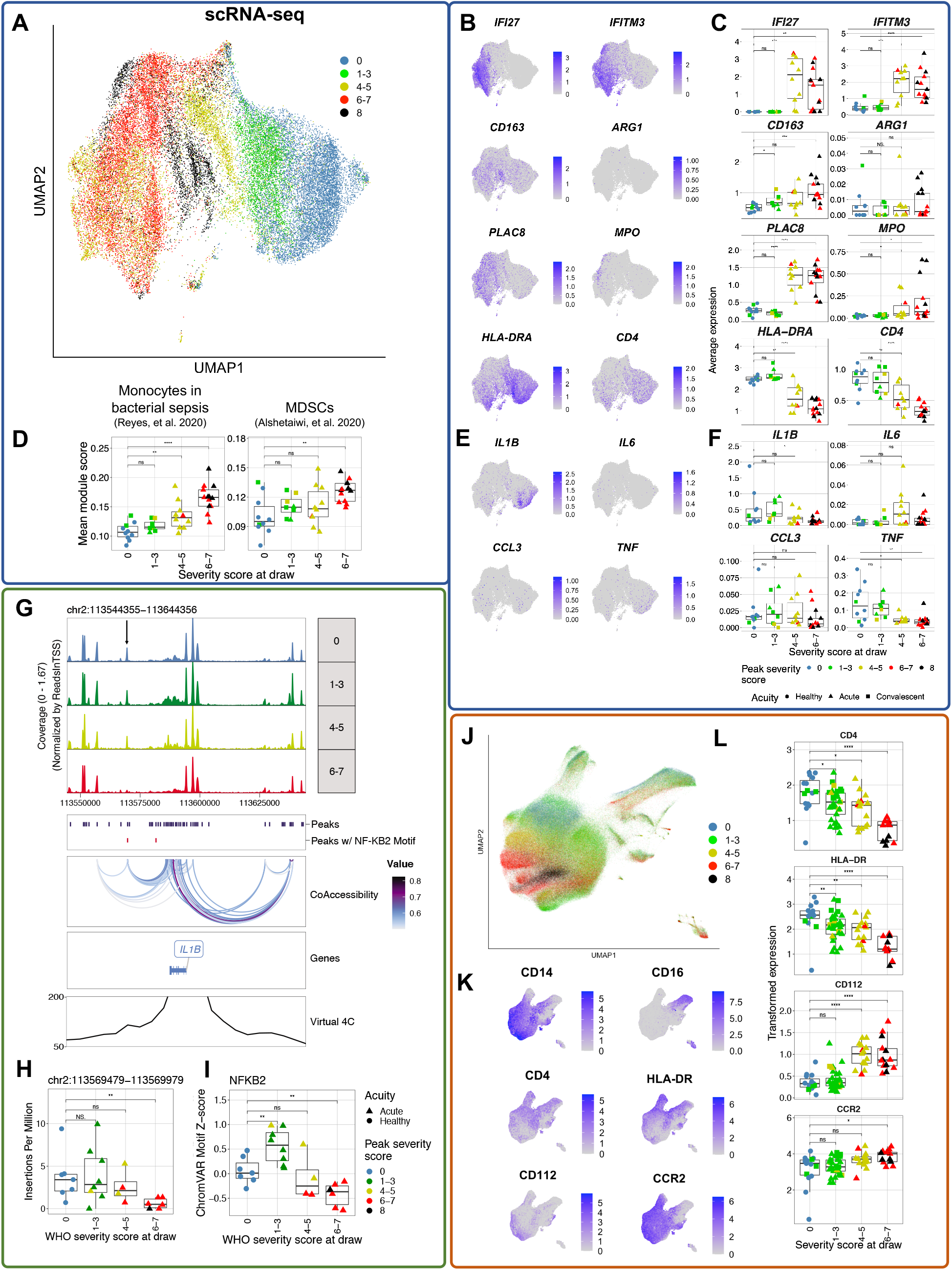
Monocytes with dysfunctional features emerge in severe and fatal COVID-19. **A)** UMAP projection of CD14 monocytes from scRNA-seq dataset, colored by peak disease severity score. **B,E)** Feature plots depicting expression of selected DEGs and pro-inflammatory cytokine-encoding genes by CD14 monocytes (see **Table S10** for complete DEG list). **C,F)** Box plots depicting average expression of features plotted in (**B,E**) in each sample. **D)** Box plot showing average module score for gene signatures of monocytes in bacterial sepsis *(43)* and MDSCs *(42)* (see **Methods**). **G)** Genome track showing 100 kb around the *IL1B* gene locus. Top panel shows the coverage at different peak regions for different severity groups; the box below shows peaks called from all CD14 monocytes in that region (dark blue) and peaks containing putative strong NF-κB2 binding sites (red); the “CoAccessibility” box shows the accessibility correlated peak pairs across all CD14 monocytes; the “Genes” box shows location of *IL1B* gene, blue color means the gene is located on the minus strand; the bottom “Virtual 4C” track shows KR-normalized contact frequencies to the *IL1B* promoter in THP1 monocytic cells. A locus with significantly altered accessibility is marked by a black arrow. **H)** Box plot depicting the accessibility of the peak marked by a black arrow in (**G**) in CD14 monocytes from each sample, measured by number of Tn5 insertions per million. **I)** Box plot depicting the average chromVAR z-scores of NF-κB2 binding motifs in CD14 monocytes from different samples. **J)** UMAP projection of all monocytes from CyTOF dataset, colored by peak disease severity score. **K-L)** Feature plots (**K**) and box plots (**L**) depicting arcsinh-transformed expression of selected protein markers by monocytes in CyTOF dataset. For all box plots: points are colored by the peak disease severity score, shaped according to disease acuity, and grouped by the disease severity score at time of sample collection. *, p < 0.05; **, p < 0.01; ***, p < 0.001; ****, p < 0.0001, n.s., not significant at p = 0.05 by two-sided Wilcoxon rank-sum test with Bonferroni’s correction for multiple hypothesis testing.

For example, we identified disease severity-associated phenotypes in cDC2 cells but not other DC subsets (**Fig. 2C-E**). In cDC2 cells, *FCER1A*, known to be involved in inflammatory DC signaling *(32)*, *CD83*, an activation marker of mature DCs *(33)*, and *CTSS*, which is involved in antigen presentation (**Fig. 2F**) *(34)* were downregulated with increased disease severity while genes associated with tolerogenic or anti-inflammatory responses, like *PKM* and *CD163*, were upregulated *(35*, *36)*. Collectively, these results indicate that dysfunctional and antiinflammatory cDC2 cells may be a feature of severe COVID-19, with important potential implications for Tfh development and mucosal immunity.

CD8 T_EM_ cells also displayed severity-associated transcriptional perturbations (**Fig. 2C, Fig. S5D-E**). Notably, several markers of CD8 effector capacity, like *PRF1, GZMB*, and *CX3CR1 (37*, *38)*, were downregulated primarily in patients with mild COVID-19 (**Fig. S5F**). Additionally, in severe and fatal COVID-19 patients, CD8 T_EM_ retained expression of markers of effector capacity without showing features of exhaustion (**Fig. S5G**). Together, these data imply that over-exuberant peripheral cytotoxic T cell responses may be associated with severe disease, similar to previous reports *(39)*.

We next examined the impact of disease time point on immune phenotype, given the heterogeneity of sampling times between patients (**Fig. 1B**). These analyses are limited by our small sample size of acutely infected mild patients in our transcriptional dataset. Nonetheless, these analyses suggest that disease acuity has more impact on transcriptional perturbation in moderate patients than in mild patients (**Fig. S6**): patients with a peak WHO severity score of 1-3 had similar transcriptional profiles marked by minimal perturbation, regardless of acuity status, while those with a peak score of 4-5 appeared to cluster by acuity. These results imply that mild COVID-19 may be marked by minimal, or rapidly resolved, systemic immune responses, a finding that is orthogonally supported by our CyTOF analyses that include a greater number of subjects. Cell types most perturbed in convalescence included CD8 T_EM_ and CD14 monocytes; in moderate patients, B cells were also perturbed in convalescence, displaying downregulation of *Ig* genes (**Fig. S6**). Additional longitudinal sampling across all severity groups is necessary to clarify these signatures.

### Emergence of monocytes with dysfunctional features in severe COVID-19

We next analyzed the phenotypes of peripheral monocytes in COVID-19, as these cells appeared to be strongly reconfigured in nonlinear dimensionality reduction projections for all three modalities (**Fig. 1C-D, F-I**). Embedding of CD14 monocytes alone from the transcriptomic dataset recapitulated this phenotypic shift (**Fig. 3A**). Similar to previous reports *(11*, *15*, *21*, *40*, *41)*, multiple ISGs (**Fig. 3B-C, Fig. S7**) and markers of immature and tolerogenic monocytes, like *CD163, PLAC8*, and *MPO* (**Fig. 3B-C**), were upregulated with increasing disease severity. Notably, *ARG1*, encoding the myeloid-derived suppressor cell (MDSC) marker and T cell inhibitor arginase, was also upregulated most prominently in the monocytes of fatal COVID-19 patients (**Fig. 3B-C**). Monocytes from severe and fatal COVID-19 patients possessed additional features of an MDSC-like phenotype, including loss of HLA class II-encoding genes (**Fig. 3B-C, Fig. S7**) and enrichment of published gene signatures from MDSCs *(42)* and monocytes in the setting of bacterial sepsis *(43)* (**Fig. 3D**). Additionally, we noted a severity-associated loss of *CD4* expression (**Fig. 3B-C**), which is involved in monocyte-to-macrophage differentiation and pro-inflammatory cytokine induction in CD14 monocytes *(44*, *45)*. These results collectively suggest that suppressive and dysfunctional monocytes are a feature of severe and fatal COVID-19, in agreement with previous reports *(11*, *15*, *21)*. Importantly, mild COVID-19 generally did not lead to this shift towards suppressive and dysfunctional monocytes.

We also noted that *C19orf59*, which encodes MCEMP1, a key biomarker for stroke risk and outcome *(46*, *47)*, was upregulated in the monocytes of severe and fatal COVID-19 patients (**Fig. S8**). Given the accumulating data that COVID-19 can drive thrombotic complications including ischemic stroke, we also examined expression of other transcripts reported to predict stroke risk by Raman, et al. *(47)*. We found that each of the five most predictive transcripts for stroke risk and prognosis reported by Raman, et al. (*C19orf59*, *IRAK3, ANXA3, RBM47*, and *TLR5*) were abundantly expressed in monocytes and neutrophils, and that each of these transcripts was significantly upregulated in severe COVID-19 in either monocytes or neutrophils (**Fig. S8**).

We next examined the expression of pro-inflammatory cytokine-encoding genes by peripheral monocytes. Similar to previous work *(11*, *15)*, we found minimal expression of key monocyte-derived pro-inflammatory cytokine-encoding genes in severe and fatal COVID-19 patients; in fact, *IL1B* and *TNF* were significantly downregulated in the monocytes of severe and fatal COVID-19 patients (**Fig. 3E-F**). Similar phenotypic changes were also observed in CD16 monocytes in the scRNA-seq dataset (**Fig. S7**). Further, monocytes from mild COVID-19 patients also did not upregulate many pro-inflammatory cytokine-encoding genes (**Fig. 3E-F**), in contrast to mild cases of similar viral infections like influenza *(48*–*50)*.

To investigate potential gene regulatory mechanisms that could explain the downregulation of *IL1B* in the CD14 monocytes of severe and fatal COVID-19 patients, we examined changes in chromatin accessibility around the *IL1B* locus. We identified a putative enhancer downstream of *IL1B*, which shows linkage to the *IL1B* promoter via single cell co-accessibility analysis and chromosome conformation capture Hi-C data from the THP-1 monocytic cell line (**Fig. 3G**, see **Methods**) *(51)*. This putative enhancer showed significantly decreased accessibility in severe COVID-19 patients (p=0.0081, Wilcoxon test, **Fig. 3H**). Furthermore, this element contains an NF-κB binding motif, suggesting it may be regulated by NF-κB family transcription factors. We identify similar changes in peaks containing NF-κB motifs at loci for other monocyte-produced inflammatory cytokines including *CCL2* and *CCL3* (**Fig. S9**).

Consistent with this hypothesis, we further observed COVID-19-associated changes in genomewide NF-κB family transcription factor activity. Using ChromVAR analysis to quantify transcription factor activity from the chromatin accessibility of each cell, we found increased NF-κB activity in mild cases and significantly decreased activity in severe cases, consistent with the changes we observed at the *IL1B* locus (p=0.0047, Wilcoxon test, **Fig. 3I**) *(52)*. Our scRNA-seq data did not show significant transcriptional changes for most NF-κB-family TFs, although *REL* and *RELB* are significantly downregulated in severe COVID-19 (**Fig. S9**). This could either reflect technical limitations of measuring lowly-expressed transcription factor transcripts, or it could indicate that our observed NF-κB activity changes are caused by post-translational modifications *(53)*. The NF-κB pathway is crucial for the inflammatory responses to virus infections in innate immune cells *(53*–*55)*, and its activation relies on various proinflammatory cytokines including IL-1β and TNFα *(56)*. Activated NF-κB can further induce *TNF* and *IL1B* expression *(53*, *57)*, leading to a positive feedback loop. Our results suggest that aberrant decreases in NF-κB activity in severe COVID-19 may result in loss of accessibility at putative enhancers of key cytokine genes, providing a potential mechanism for the previously reported impairment in cytokine production by peripheral monocytes in COVID-19 *(11)*.

We also examined the epigenetic regulation at the *CD4* locus, as this gene was significantly downregulated with increasing disease severity. Although there was no change in chromatin accessibility of the *CD4* gene promoter between severity groups, we found that the accessibility of monocyte-specific *CD4* gene putative regulatory regions was significantly reduced in severe samples (**Fig. S9**). This monocyte-specific loss of CD4 expression may provide an additional mechanism explaining the previously reported impairment of cytokine production by monocytes in COVID-19 *(11*, *21)*, as the interaction between IL-16 and monocytic CD4 induces the expression of proinflammatory cytokines, including IL-1β *(44)*.

Profiling of COVID-19 monocytes by CyTOF recapitulated many of these findings (**Fig. 3J,K**). This included a loss of CD16^+^ monocytes as well as a distinct shift in the phenotype of CD14^+^ monocytes (**Fig. 3J,K; Fig. S10**). As observed in our transcriptional data, expression of HLA-DR and CD4 was lost in the CD14^+^ monocytes of severe COVID-19 samples. Importantly, this proteomic reconfiguration was not observed in patients with mild COVID-19, evident in nonlinear dimensionality reduction (**Fig. 3J**). Mild patients experienced no significant increase in expression of the stress marker CD112, nor did they upregulate CCR2, which is involved in monocyte recruitment to the airways in the setting of severe COVID-19, although both of these markers were upregulated in severe patients (**Fig. 3J-L**) *(58*, *59)*. They also displayed a less dramatic loss of HLA-DR and CD4 expression compared to monocytes in severe cases (**Fig. 3J-L**). These results indicate that while monocytes are dramatically remodeled in severe COVID-19, mild COVID-19 has minimal, or rapidly resolved, impact on the monocyte proteome.

### Peripheral NK cells are depleted in severe COVID-19 and have a highly activated phenotype

We next interrogated changes in the NK cells of COVID-19 samples. As demonstrated previously *(15*, *60)*, peripheral NK cells were substantially depleted across all three modalities and transcriptionally reconfigured in severe COVID-19 samples (**Fig. 4A-B, Fig. S11**), which may reflect their trafficking to the site of infection. Transcriptional changes appeared to be driven by upregulation of several canonical NK cell activation genes, including higher expression of cytotoxic effector molecule-encoding genes *GZMB* and *PRF1*, as well as proliferation marker *MKI67* and ISGs like *XAF1* (**Fig. 4B-C**). NK cells from moderate and severe, but not mild, COVID-19 cases, also displayed transcriptional evidence of exhaustion (**Fig. 4C**).

**Fig. 4.**
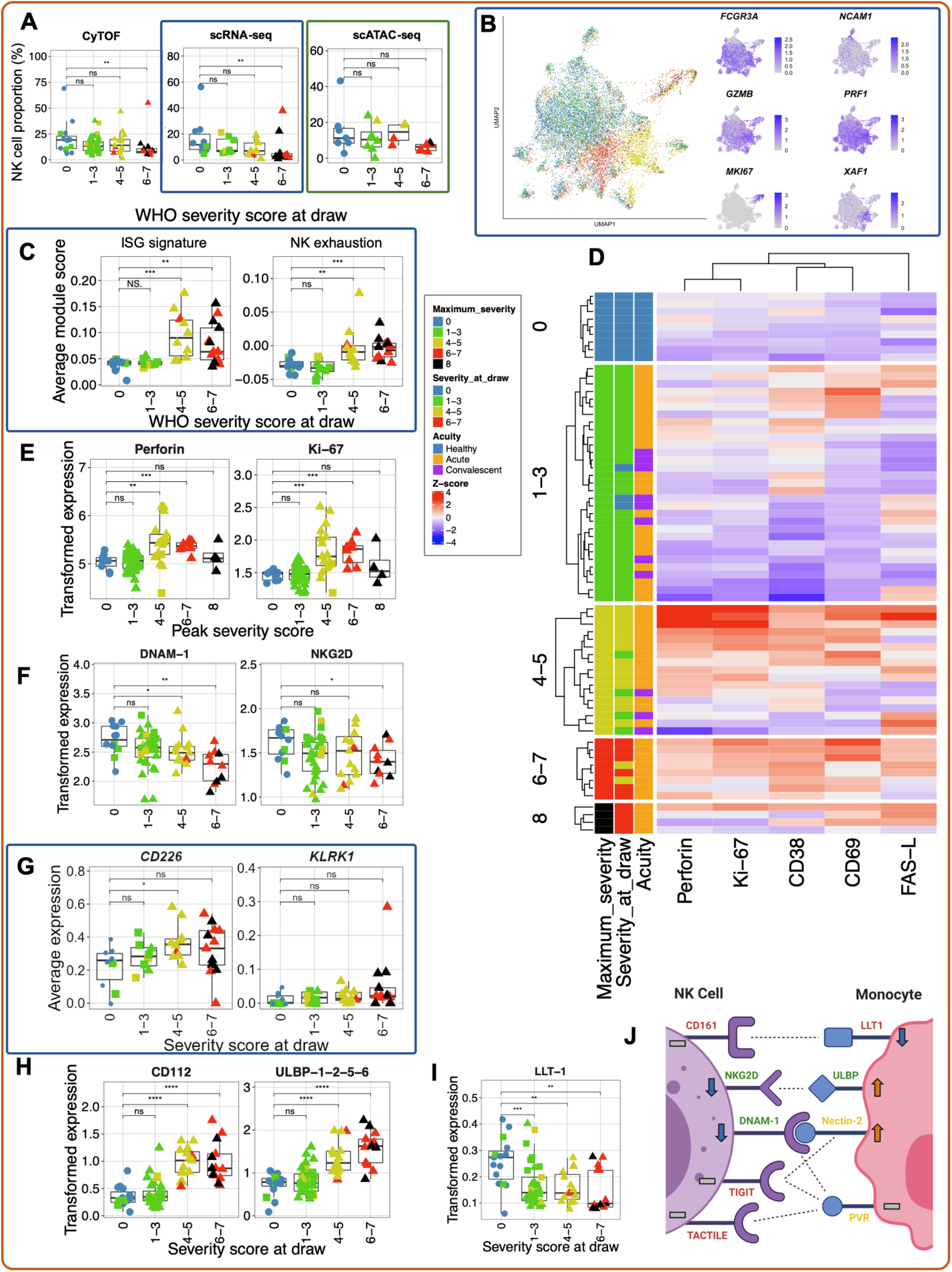
NK cells of severe COVID-19 patients exhibit a unique proteomic and transcriptional profile. **A)** Box plots of manually-annotated NK cell proportions from CyTOF (left), Seurat v4-annotated NK cell proportions from scRNA-seq (center), and Seurat v4-annotated NK cell proportions from scATAC-seq (right, see **Methods**). **B)** UMAP projections of NK cells from scRNA-seq dataset colored by peak disease severity score (left) and selected DEGs (right; see **Table S11** for complete list). **C)** Box plots of average ISG signature and NK cell exhaustion (defined as expression of *LAG3, PDCD1* and *HAVCR2*, see **Methods**) module scores in Seurat v4-annotated NK cells. **D)** Heatmap of Z-score normalized protein-level expression of canonical NK cell activation and cytotoxicity markers (Perforin, Ki-67, CD38, CD69, and Fas-L) in each sample. **E)** Box plots quantifying arcsinh-transformed average expression of Ki-67 (left) and Perforin (right) by NK cells, grouped by peak disease severity score. **F)** Box plots of average expression of DNAM-1 (left) and NKG2D (right) protein in activated NK cells (defined as CD38^+^CD69^+^ NK cells based on manual gating) from CyTOF dataset. **G**) Box plots of average expression of *CD226* (which encodes DNAM-1; left) and *KLRK1* (which encodes NKG2D; right) from scRNA-seq dataset (**G**). **H-I)** Box plots of arcsinh-transformed average protein-level expression of NK cell ligands CD112 and ULBP-1,2,5,6 (**H**) and LLT-1 (**I**) by monocytes. **J)** Schematic illustrating changes in proteinlevel expression of NK cell activating and inhibitory receptors and their ligands. Text color indicates if a receptor/ligand is activating (green), inhibitory (red), or either, depending on the context (yellow). Arrows and dashes indicate whether abundance of a protein is increased, decreased, or unchanged in severe COVID-19 compared to healthy controls. Dashed lines indicate interactions between receptors and ligands. For all box plots except (**E**): points are colored by peak disease severity score, shaped according to disease acuity, and grouped by the severity score at sample collection. *, p < 0.05; **, p < 0.01; ***, p < 0.001; ****, p < 0.0001, n.s., not significant at p = 0.05 by two-sided Wilcoxon rank-sum test with Bonferroni’s correction for multiple hypothesis testing.

We next examined this NK cell activation signature at the protein level. We corroborate previously known changes in NK cell biology, including increased expression of the activation markers CD38 and CD69 *(60)*, and also demonstrate that surface expression of the death receptor ligand Fas-L is increased in moderate and severe COVID-19 patients (**Fig. 4D**; **Fig. S11**). While perforin was also upregulated in moderately and severely ill patients, surprisingly, NK cells from fatal COVID-19 patients failed to upregulate both this cytotoxic effector and the proliferation marker Ki-67 (**Fig. 4D-E**). These data, coupled with transcriptomic evidence of NK cell exhaustion in severe and fatal COVID-19 (**Fig. 4C**), suggest that defects in NK cell cytotoxicity may be associated with adverse outcomes.

### Dynamic changes in NK cell receptors and ligands may underlie COVID-19 severity-associated NK activation

To assess mechanisms of NK cell activation, we interrogated changes in the NK cell repertoire of surface-expressed activating and inhibitory receptors as well as their cognate ligands. Notably, surface expression of the activating receptors DNAM-1 (CD226) and NKG2D was significantly downregulated in the activated (CD69^+^CD38^+^) (**Fig. 4F**) and total (**Fig. S11F**) NK cells of severe COVID-19 samples compared to healthy controls, despite no change in the expression of the genes encoding these proteins in the total NK cells within our scRNA-seq data (**Fig. 4G**). As expression of both DNAM-1 and NKG2D can be downmodulated following ligation *(61*, *62)*, we investigated the abundance of the two ligands for DNAM-1, poliovirus receptor (CD155) and Nectin-2 (CD112), and of the ULBP proteins, which are recognized by NKG2D. Both Nectin-2 and the ULBPs were significantly upregulated on the peripheral monocytes of hospitalized COVID-19 patients compared to healthy controls (**Fig. 4H**), which supports the hypothesis that SARS-CoV-2 may decrease surface expression of DNAM-1 and NKG2D through internalization following ligation of overexpressed Nectin-2 and ULBP proteins due to stress. We found no change in the expression of the other ligand for DNAM-1, CD155, nor did we observe any changes in the protein- or RNA-level expression of either TIGIT, an inhibitory receptor that competes with DNAM-1 for binding of Nectin-2, or TACTILE (CD96), which recognizes CD155 (**Fig. S11E**). Alternatively, activated NK cells expressing DNAM-1 or NKG2D may migrate to the tissue in the setting of severe disease, depleting these markers from the circulating population.

We also investigated the surface expression levels of other ligands for NK cell receptors and observed a loss of LLT-1 expression on CD14^+^ monocytes that appears to correlate with disease severity, with a near-total loss in fatal samples (**Fig. 4I**); however, we found no change in the expression of the inhibitory receptor that recognizes LLT-1, CD161, on NK cells (**Fig. S11G**). The overall profile of activating and inhibitory receptors and ligands expressed in severe COVID-19 is summarized in **Fig. 4J** and suggests that the activated phenotype observed in these samples may be driven by activating signals received through DNAM-1 and NKG2D as well as a lack of inhibitory signaling through CD161.

### A hyperactivated neutrophil signature marks severe and fatal COVID-19

Despite evidence that neutrophils are major players in the dysregulated immune response defining severe and fatal COVID-19 *(16*, *18*–*20*, *63*–*66)*, there is a relative dearth of deep profiling of neutrophils from COVID-19 patients given their sensitivity to both cryopreservation and mechanical stimulation *(67*, *68)*. To address this, we demonstrated that Seq-Well generated high-quality scRNA-seq data of primary human neutrophils from a healthy blood donor (**Fig. S12**). While fewer genes were detected in sequenced neutrophils, the number of unique molecular identifiers (UMIs) sequenced in neutrophils was comparable to the expected recovery based on known RNA content *(69*, *70)*. We also found that neutrophils from ACK-lysed whole blood were phenotypically similar to neutrophils isolated by magnetic-activated cell sorting (MACS; **Fig. S12D-E**); the former strategy was also able to uncover other granulocytic cell types like eosinophils (**Fig. S12C**).

Seq-Well processing of red blood cell-lysed whole blood yielded 33,276 high-quality single transcriptomes of primary neutrophils (**Fig. 5A**). These cells uniformly and specifically expressed neutrophil lineage marker-encoding genes, including *CSF3R* and *CXCR2*, indicating their identity as canonical neutrophils (**Fig. 5A**, **Table S6**). Non-linear dimensionality reduction revealed that neutrophil transcriptional phenotype was strongly remodeled in COVID-19 (**Fig. 5A**), driven in part by upregulation of *PADI4*, which is required for NETosis, the IL-8 receptor *CXCR1*, and multiple alarmins, including *S100A8* and *S100A9* (**Fig. 5B-C**), which together induce neutrophil chemotaxis and adhesion *(71)*. We also noted disease severity-specific induction of interferon-stimulated genes (ISGs) in moderate and severe, but not in mild, COVID-19 patients (**Fig. 5D**). While this ISG signature was detected across most cell types in moderately and severely ill patients, neutrophils upregulated the broadest number of ISGs (**Fig. S13**). Importantly, the differential expression of ISGs by neutrophils between COVID-19 severity groups was not due to a difference in infection time point between patients: neutrophils from patients with mild COVID-19 did not upregulate ISGs at any point during infection (**Fig. 5D**). Additionally, gene set enrichment analysis demonstrated upregulation of genes associated with neutrophil phagocytosis and degranulation in a disease severity-associated fashion (**Fig. 5E**, see **Table S12**).

**Fig. 5.**
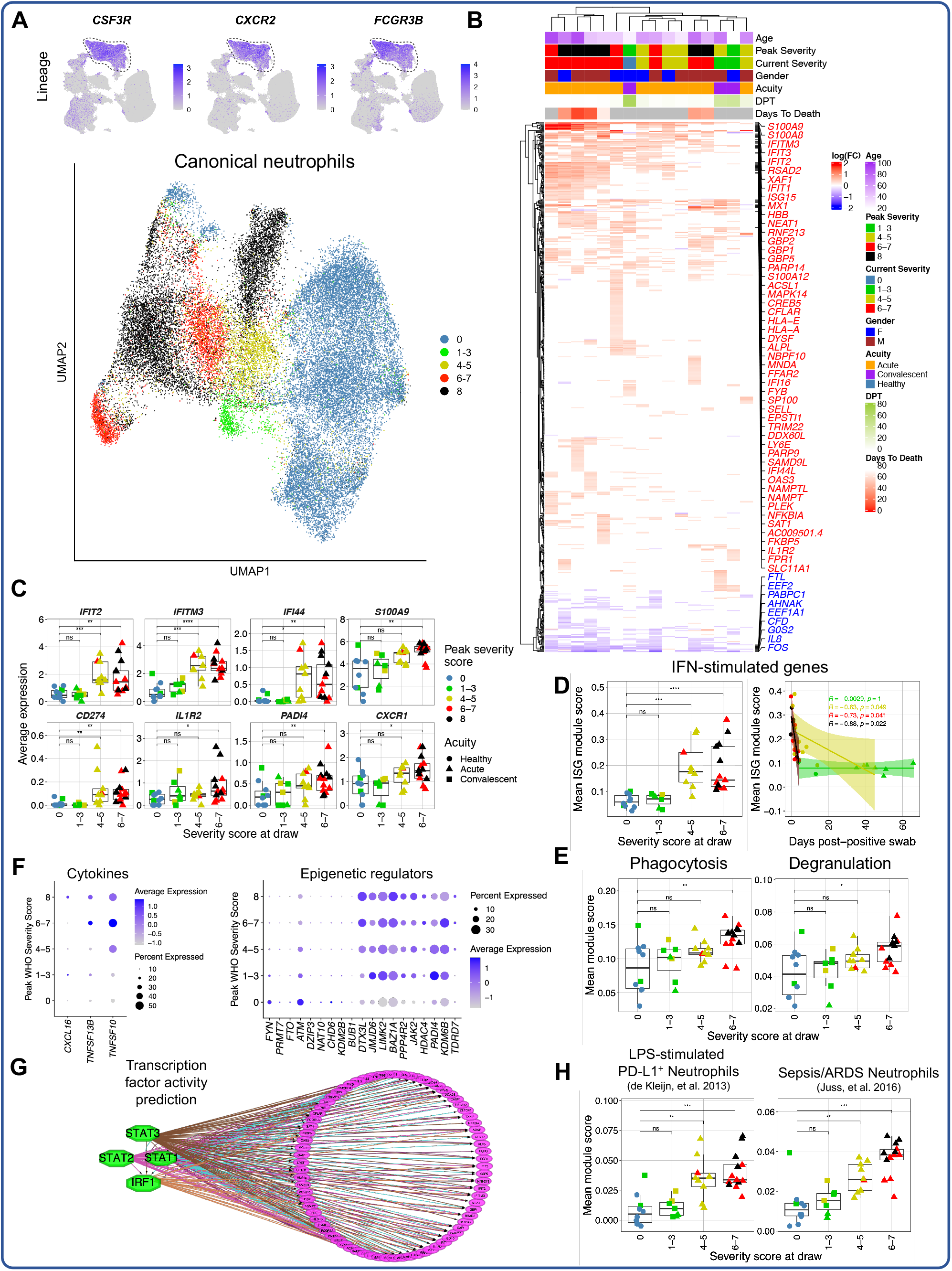
Neutrophil activation is a hallmark of severe and fatal COVID-19. **A)** UMAP projections of complete scRNA-seq dataset colored by expression of canonical neutrophil markers (top) and of canonical neutrophils alone colored by peak disease severity score (bottom). **B)** Heatmap of DEGs between neutrophils of each COVID-19 sample compared to neutrophils of all healthy controls, colored by average log(fold-change). All displayed DEGs are statistically significant at the p < 0.05 confidence level by Seurat’s implementation of the Wilcoxon rank-sum test (two-sided, adjusted for multiple comparisons using Bonferroni’s correction). **C)** Box plots depicting average expression of selected neutrophil DEGs by severity group (see **Table S13** for complete DEG list). **D)** Plots depicting median ISG signature score of neutrophils in each sample grouped by disease severity score at time of sample collection (left) and by days after first positive NP swab (right). All points are colored by peak disease severity score. For scatter plot at right, Pearson’s r, exact two-sided p values and the 95% confidence interval are shown for each peak disease severity score grouping. **E)** Box plots depicting average module scores for genes sets of neutrophil phagocytosis and neutrophil degranulation (see **Methods**, **Table S12**). **F)** Dot plots depicting average and percent expression of pro-inflammatory cytokine encoding genes (left) and epigenetic regulators (right) by canonical neutrophils. The *y* axis corresponds to the peak disease severity score. **G)** Results of transcription factor activity prediction analysis performed by iRegulon *(76)*. DEGs between neutrophils from severely ill (peak severity 6-8) and neutrophils from healthy controls were used as input (see **Methods, Table S14**). **H)** Box plots of average module scores for PD-L1^+^ neutrophils in an *in vitro* model of endotoxemia *(74)* and granulocytes in the setting of sepsis and ARDS *(75)* (see **Methods, Table S12**). For all box plots: points are colored by the peak disease severity score, shaped according to disease acuity, and grouped by the disease severity score at time of sample collection. *, p < 0.05; **, p < 0.01; ***, p < 0.001; ****, p < 0.0001, n.s., not significant at p = 0.05 by two-sided Wilcoxon rank-sum test with Bonferroni’s correction for multiple hypothesis testing.

We also identified two distinct neutrophil immunophenotypes of fatal COVID-19. Neutrophils from 4/6 fatal COVID-19 cases had robust ISG induction and expressed *CD274* (encoding PD-L1; **Fig. 5B-C**), in line with previous work *(21)*. However, we also identified 2 fatal COVID-19 cases with less pronounced ISG induction but with upregulation of additional neutrophil activation markers not observed in other samples, including *CXCR4*, *CLEC12A*, *EGR1*, and the decoy IL1β receptor *IL1R2* (**Fig. 5B-C**). Additional severity-associated changes in neutrophil phenotype included the upregulation of pro-inflammatory cytokine-encoding genes, including *CXCL16* and *TNFSF10* (encoding TRAIL), as well as upregulation of several epigenetic regulators involved in neutrophil activation, like *PADI4*, which is required for formation of neutrophil extracellular traps (NETs; **Fig. 5F**) *(19*, *72*, *73)*.

Transcription factor activity analysis implicated STAT1, STAT2, STAT3, and IRF1 as key upstream regulators of the observed transcriptional reconfiguration, further suggesting that COVID-19 neutrophils are strongly activated by type I IFN in a disease severity-specific fashion (**Fig. 5G**). To better contextualize these findings, we scored the neutrophils in our dataset against gene modules upregulated in a model of endotoxemia *(74)* and in ARDS-complicated sepsis *(75)*. This analysis revealed that both of these signatures are upregulated with increasing COVID-19 severity (**Fig. 5H**), implying similarities in neutrophil phenotype between these clinical conditions. Collectively, profiling fresh whole blood rather than isolated PBMCs reveals a prominent neutrophil hyperactivation signature in severe and fatal COVID-19.

### Developing neutrophils are a feature of fatal COVID-19

We next analyzed a population of developing neutrophils from the transcriptomic dataset that was enriched in patients with severe and fatal COVID-19 (**Fig. 1E, Fig. 6A**). This population has been identified in other COVID-19 datasets but is not yet well-characterized *(21*, *63)*. These cells specifically and highly expressed genes encoding markers expressed at distinct stages in neutrophil development, including *DEFA1B, LCN2*, and *MMP8* (**Fig. S1C, Table S6**), indicating that they likely represent immature neutrophils derived from emergency granulopoiesis. As we hypothesized that these cells were not a static population but rather were dynamically differentiating, we embedded them in two and three dimensions using PHATE, a dimensionality reduction method developed to visualize phenotypic continua, branches, and continual progressions (**Fig. 6B-C, Fig. S14**) *(77)*. Clustering of these cells revealed 5 clusters that corresponded to sequential stages in neutrophil development, beginning with cluster 0 (pro-neutrophils) expressing primary neutrophil granule protein-encoding genes, followed by clusters 1-3 (pre-neutrophils) consecutively expressing secondary and tertiary neutrophil granule protein-encoding genes, and finally cluster 4 (mature neutrophils) which expresses markers of canonical neutrophils (**Fig. 6C-D**) *(78)*. Importantly, *ELANE* (which encodes neutrophil elastase), was specifically expressed by pro-neutrophils, implying that these cells may be capable of NETosis *(79*, *80)*. An orthogonal approach ordering each cell in latent time modeled by splicing kinetics of RNA velocity *(81*, *82)* (**Fig. 6E**) also revealed a similar developmental trajectory with respect to both granule protein-encoding genes (**Fig. 6F**) and transcription factors known to be involved in neutrophil development, like the C/EBP family (**Fig. 6G, Fig. S14**). In our earlier work, we hypothesized that developing neutrophils may arise via transdifferentiation from plasmablasts based on their phenotypic similarity in nonlinear dimensionality reduction manifold space and subsequent analysis of cellular trajectory by RNA velocity. In this larger dataset, a phenotypic relationship between developing neutrophils and plasmablasts was still evident (**Fig. 1C-D**), but RNA velocity analysis of plasmablasts, developing neutrophils, and mature neutrophils did not reveal a clear transdifferentiation bridge (**Fig. S14**). Orthogonal experiments are necessary to conclusively determine the development origins of these cells.

**Fig. 6.**
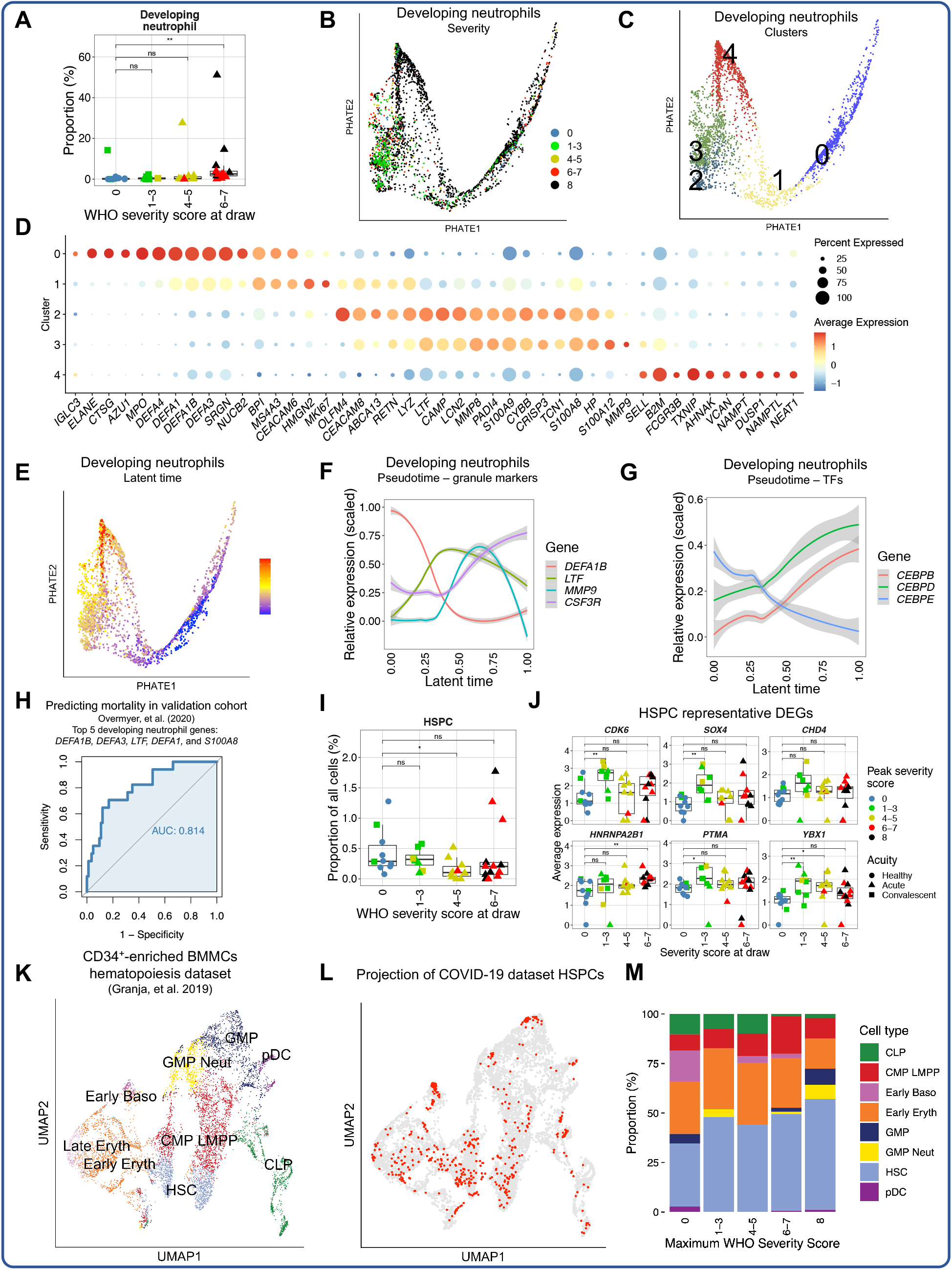
Emergency granulopoiesis and other hematopoietic abnormalities characterize severe and fatal COVID-19. **A)** Box plot depicting proportions of developing neutrophils in each sample from the scRNA-seq dataset. **B-C)** Twodimensional PHATE projection of developing neutrophils colored by peak disease severity score (left) and cluster number (right). **D)** Dot plot depicting percent and average expression of DEGs between developing neutrophil clusters (see **Table S15**). **E)** Twodimensional PHATE projection of developing neutrophils colored by latent time calculated by scVelo *(82)*. **F-G)** Scaled expression of selected neutrophil granuleencoding genes (**F**) and C/EBP transcription factor family-encoding genes (**G**) by developing neutrophils across inferred latent time. **H)** Receiver operating characteristic (ROC) curve depicting sensitivity and specificity of 28-day mortality prediction of a five-gene signature of developing neutrophils (*DEFA1B, DEFA3, LTF, DEFA1*, and *S100A8*) in an independent validation cohort of 103 samples where 17 cases are fatal *(91)*. **I)** Box plot depicting proportions of Seurat v4-annotated HSPCs in scRNA-seq dataset. **J)** Box plots of average expression of selected HSPC DEGs (see **Table S16**). **K-L)** UMAP projection of Seurat v4-annotated HSPCs from scRNA-seq dataset into a publicly-available blood and bone marrow hematopoiesis dataset *(90)* colored by published cell type annotations (**K**) and with projected HSPCs colored in red (**L**). **M)** Bar plot depicting proportions of cell type identities transferred after projection into the publicly-available hematopoiesis dataset for each peak disease severity score bin. For all box plots: points are colored by the peak disease severity score, shaped according to disease acuity, and grouped by the disease severity score at time of sample collection. *, p < 0.05; **, p < 0.01; ***, p < 0.001; ****, p < 0.0001, n.s., not significant at p = 0.05 by two-sided Wilcoxon rank-sum test with Bonferroni’s correction for multiple hypothesis testing.

As developing neutrophils were present uniformly, often at high frequencies, in patients with fatal COVID-19, and these cells specifically expressed many genes not found in other peripheral blood cell types, we hypothesized that the developing neutrophil gene signature might be an accurate predictor of mortality in COVID-19. We therefore identified the five most positively differentially-expressed genes between developing neutrophils (6,569 cells across 21 patients) and all other cells in our dataset: *DEFA1B, DEFA3, LTF, DEFA1*, and *S100A8*. We next obtained a publicly-available whole blood bulk transcriptomic dataset of 103 COVID-19 patients as a validation cohort and scored each sample in this dataset by the aggregated expression of these five genes (see **Methods**). After scoring each sample, we used the associated patient metadata to determine the 28-day mortality of each scored sample. We then constructed a receiver operating characteristic (ROC) plot using the gene score as predictor and the 28-day mortality as the response variable. We found that the developing neutrophil gene score accurately predicted 28-day mortality of the 17 patients who succumbed to infection (AUC = 0.81; **Fig. 6H, Fig. S14**). Importantly, the sequential organ failure assessment (SOFA) score at the time of sample collection did not strongly predict 28-day mortality (AUC = 0.67), indicating that the presence of developing neutrophils is a better prognostic indicator than current clinical status measured by clinically used severity scales (**Fig. S14**). Thus, developing neutrophils are likely enriched in the blood of fatal COVID-19 cases in other cohorts, and gene signatures from these cells have promise as a potential prognostic indicator.

### Myeloid skewing of hematopoietic progenitors in severe and fatal COVID-19

Considering that severe and fatal COVID-19 patients displayed evidence of emergency myelopoiesis in the periphery, we hypothesized that there also may be severity-associated aberrations in a small population of hematopoietic stem and progenitor cells (HSPCs) we identified in our transcriptomic dataset (**Fig. 1C-D, Fig. 2C**). Although the frequency of these cells did not change in COVID-19 (**Fig. 6I**), we found that several genes involved in HSPC maintenance and self-renewal, including *CDK6, SOX4*, and *CHD4 (83–89*), were generally upregulated in COVID-19 patients relative to healthy controls (**Fig. 6J**). These DEGs suggest that the hematopoietic progenitor compartment in COVID-19 patients has been transcriptionally reshaped to accommodate the stress of emergency hematopoiesis. To better understand the identities of these cells, we leveraged a publicly available single-cell transcriptomic dataset of hematopoiesis in healthy human blood and bone marrow (**Fig. 6K**) *(90)*, into which we projected HSPCs from our COVID-19 dataset (**Fig. 6L**). We found a trend towards myeloid skewing in COVID-19 circulating HSPCs with increasing disease severity, with lower frequencies of common lymphoid progenitors in severe and fatal patients, as well as granulocyte/macrophage progenitor (GMP)/neutrophil-like cells appearing in severe and fatal cases (**Fig. 6M**). Together, these results indicate severity-associated changes in hematopoiesis in COVID-19 with greater myeloid skewing evident in circulating HSPCs.

## DISCUSSION

In this work, we have compiled a trimodal single-cell atlas of immune cells from COVID-19 patients with a wide range of disease severities through scRNA-seq, scATAC-seq, and CyTOF. By virtue of our whole blood analyses and multi-modal approach, our analysis reveals novel mechanisms of immune activation and dysregulation in the setting of severe COVID-19, in addition to providing critical validation of results from other studies. We found that neutrophils and NK cells appeared strongly activated with increasing disease severity, with heightened ISG induction and increased expression of cytotoxic machinery. Conversely, monocytes and dendritic cells displayed dysregulated and tolerogenic features in severe and fatal COVID-19. Finally, we found dramatic changes in hematopoietic development in severe COVID-19, with the appearance of a population of developing neutrophils in the periphery and skewing of circulating hematopoietic precursors towards the myeloid lineage.

Importantly, our profiling of patients with mild COVID-19 allows us to demonstrate that many of these changes occur largely in severe and fatal COVID-19 and not in milder forms of the disease. For example, the kinetics and role of local and systemic IFN signaling in ameloriating or exacerbating SARS-CoV-2 remain controversial *(41*, *92*–*95)*. Here, we noted minimal induction of ISGs in mild COVID-19 cases, regardless of the time point of infection. This suggests that robust IFN responses detectable in the periphery may not be required for disease resolution. Additionally, we observed surprisingly little perturbation of monocytes and NK cells from mild COVID-19 patients, whereas mild cases of influenza are known to induce systemic activation of these cells *(96*, *97)*. Direct comparative analyses and larger sample sizes will be necessary to identify conserved or differential features between mild COVID-19 and other mild respiratory virus infections.

Our analysis demonstrated a strong severity-associated hyperactivation phenotype in peripheral neutrophils marked by broad ISG induction, pro-inflammatory cytokine-encoding gene production, enrichment of phagocytosis and degranulation gene sets, and upregulation of epigenetic regulators associated with inflammatory neutrophils, such as *PADI4* which is required for NETosis. Neutrophils in severe COVID-19 also strongly upregulated *S100A8* and *S100A9*, which dimerize to form the inflammatory molecule calprotectin, which is involved in neutrophil activation and chemotaxis *(71)*. Additionally, we found features associated with neutrophil exhaustion, like upregulation of *CD274*, similar to other studies *(21)*. While this finding has led some groups to conclude that neutrophils become “suppressive” in severe COVID-19 *(21)*, we believe our findings, combined with accumulating evidence that NETosis contributes to tissue injury and thrombotic complications in severe COVID-19 *(16*, *18*, *64*–*66)*, suggest a predominantly pathogenic role for circulating neutrophils in severe COVID-19. Importantly, these data provide insight into the mechanistic pathways that drive neutrophil activation. Targeting such pathways may provide new therapeutic opportunities. For example, an agonist of a neutrophil-expressed inhibitory receptor Siglec-10 (SACCOVID) has shown promising results in suppressing hyperinflammation in severe COVID-19 and is in a phase III clinical trial *(98*–*100)*. Agonists against other neutrophil-expressed inhibitory receptors, like Siglec-9, may also represent novel therapeutic candidates *(101)*.

In addition to identifying features of severe COVID-19, we were also able to identify key differences between the immune responses of patients with severe COVID-19 who went on to survive the disease versus those who did not. A striking difference between fatal and non-fatal cases was the emergence of a population of developing neutrophils that was first described by our group *(15)*. In this earlier work we identified these cells in 4/4 patients with acute respiratory distress syndrome (ARDS) requiring mechanical ventilation, including in one patient who succumbed to infection. We now show the presence of these cells in 17 additional patients, including 5/7 patients with severe and 6/6 patients with fatal COVID-19. Our trajectory analyses demonstrate that these cells follow the stages of canonical neutrophil development, beginning with defensin-rich promyelocytes and differentiating through metamyelocytes and bands to form mature neutrophils. This process may be driven by elevated levels of circulating pro-inflammatory cytokines, like IL-17, that may induce formation of neutrophils *(102)*. Although any functional or pathological role for these cells in COVID-19 pathogenesis remains unclear, their abundance in the periphery of patients with fatal COVID-19 enabled us to demonstrate that their most defining transcripts could be used to predict 28-day mortality in an independent bulk transcriptomic dataset. While emergency myelopoiesis is known to be a feature of bacterial sepsis *(103*, *104)*, it is likely that emergency myelopoiesis is also an underappreciated feature of severe viral infection. For example, in an integrated multicohort analysis of viral disease severity, Zheng, et al. report a gene module that includes several markers of immature neutrophils (eg. *CEACAM8, DEFA4, LCN2*) that is enriched in other viral infections including influenza and RSV *(105)*. Additionally, a six-mRNA classifier of viral disease severity, which includes *DEFA4*, trained on non-COVID-19 viral infections was found to also predict COVID-19 severity *(106)*. These results suggest that emergency myelopoiesis is a common feature of multiple severe viral infections and may be used to predict adverse outcomes.

In addition to canonical and immature neutrophils, the phenotype of NK cells in fatal COVID-19 cases was also distinct from that of severe non-fatal cases. While circulating NK cells are known to become activated in severe COVID-19 *(60)*, our work is the first to show that patients with fatal COVID-19 fail to proliferate and upregulate the cytotoxic effector perforin. The absence of this activation could indicate a defect in the functional responses of NK cells in fatal COVID-19, although this finding requires direct validation. We also identify NK cell ligand-receptor axes that may contribute to their activation in severe COVID-19; for example, our data suggest that DNAM-1-mediated recognition of Nectin-2 could drive NK cell activation in COVID-19, as could NKG2D-mediated activation. These data are consistent with other viral infections in which activation of NK cells through DNAM-1 or NKG2D are important *(107*, *108)*, and highlight pathways that should be investigated through *in vivo* model systems for their role in disease outcome.

While neutrophils and NK cells are activated in COVID-19, monocytes and dendritic cells appear to take on an impaired inflammatory phenotype. Perturbations in the cDC2 dendritic cells of patients with severe COVID-19 could inhibit priming of Tfh cells and development of mucosal immunity *(109)*. Both cDC2s and monocytes of critically ill COVID-19 patients appeared to downregulate or failed to upregulate genes involved in activation and inflammation; for example, cDC2s lost expression of *FCER1A* and *CD83*, while CD14^+^ monocytes notably did not upregulate any genes encoding proinflammatory cytokines such as IL-6 and CCL3, and genes encoding IL-1β and TNF were significantly downregulated in these cells compared to healthy controls. This lack of proinflammatory cytokine expression is of note as other viral infections such as influenza drive an increase in the production of these molecules by peripheral monocytes *(97)*. We identified the inactivation of NF-κB family TFs as a possible epigenetic mechanism for this silencing of proinflammatory cytokine production in monocytes, as we observed a striking loss of accessibility at an NF-κB binding site within the *IL1B* locus. This observed inactivation is unusual, given that these TFs typically play an important role in antiviral immune responses, including through regulating the production of cytokines *(110)*. Indeed, NF-κB family TFs have been implicated in driving the proinflammatory cytokine production in other cell types during SARS-CoV-2 infection *(111)*. It is also possible that the absence of NF-κB activity and accessibility at cytokine-encoding genes reflects the immaturity of monocytes in severe COVID-19.

There are several limitations to this work that should be noted. Though large for a multi-modal dataset, our sample size is still limited. Moreover, we were unable to profile each patient by all three single-cell modalities, preventing us from performing cross-modality validation on a per-patient basis. Additionally, our CyTOF panels only allowed us to examine a limited number of cell types, preventing us from performing orthogonal validation of some transcriptional or epigenetic findings in proteomic space. This work profiled exclusively circulating immune cells; while understanding the peripheral immune system is critical to understanding aberrant and protective immune responses to SARS-CoV-2 infection, it does not capture the immune response at the site of infection. Finally, further functional experimentation is necessary to validate or refute many of the hypotheses presented here.

Collectively, our work represents the first trimodal epigenomic, proteomic, and transcriptomic cell atlas of peripheral immune responses to COVID-19 across a broad spectrum of disease severity. By identifying novel immune features associated with COVID-19 mortality, as well as the immune status of patients with mild disease, our work enhances our understanding of pathological vs. protective immune responses and highlights several opportunities for therapeutic development.

## Acknowledgments

We are grateful to all participants in this study. We thank Sopheak Sim and the Stanford Functional Genomics Facility for the use of the Fragment Analyzer, as well as Angela Detweiler and Michelle Tan at the Chan Zuckerberg Biohub for assistance with sequencing. We thank Anne-Maud Ferreira for insightful conversation on mass cytometry analysis. We thank Yuhan Hao and Rahul Satija for helpful discussions on the calculation and interpretation of the perturbation score. We thank Martin Prlic for assistance with mass cytometry gating schemes. We thank Nima Aghaeepour for his assistance in implementing the *CytoNorm* package.

## Stanford COVID-19 Biobank

Thanmayi Ranganath, Nancy Q. Zhao, Aaron J. Wilk, Rosemary Vergara, Julia L. McKechnie, Lauren de la Parte, Kathleen Whittle Dantzler, Maureen Ty, Nimish Kathale, Giovanny J. Martinez-Colon, Arjun Rustagi, Geoff Ivison, Ruoxi Pi, Madeline J. Lee, Rachel Brewer, Taylor Hollis, Andrea Baird, Michele Ugur, Michal Tal, Drina Bogusch, Georgie Nahass, Kazim Haider, Kim Quyen Thi Tran, Laura Simpson, Hena Din, Jonasel Roque, Rosen Mann, Iris Chang, Evan Do, Andrea Fernandes, Shu-Chen Lyu, Wenming Zhang, Monali Manohar, James Krempski, Rosen Mann, Anita Visweswaran, Elizabeth J. Zudock, Kathryn Jee, Komal Kumar, Jennifer A. Newberry, James V. Quinn, Donald Schreiber, Euan A. Ashley, Catherine A. Blish, Andra L. Blomkalns, Kari C. Nadeau, Ruth O’Hara, Angela J. Rogers, Samuel Yang.

## Author contributions

A.J.W., M.J.L., B.W., B.P., W.J.G., and C.A.B conceived the project and designed experiments. A.L.B., R.O., E.A., K.C.N., S.Y., A.J.R., and C.A.B. conceived the clinical cohort, obtained clinical samples, and provided clinical input. A.J.W., D.J-M., E.A., and A.J.R. obtained metadata for clinical samples. T.R. coordinated clinical sample processing. Stanford COVID-19 biobank enrolled, consented, and processed clinical samples. A.J.W., M.J.L., R.P., and N.Q.Z. acquired single-cell transcriptomic data. B.W. and W.B. acquired scATAC-seq data. M.J.L., R.P., G.J.M-C., and T.R. acquired mass cytometry data. A.J.W., M.J.L., B.W., and B.P. performed bioinformatic and statistical analyses. S.T., S.H., and M.R. provided intellectual input. A.J.W., M.J.L., B.W., B.P., W.J.G., and C.A.B. wrote the manuscript, which was reviewed by all authors.

## Data and materials availability

Processed matrices (scRNA-seq and scATAC-seq) or fcs files (mass cytometry) with deidentified metadata supporting this publication are available at ImmPort (https://www.immport.org) under study accession SDY1708. Data from scRNA-seq and scATAC-seq will be deposited to GEO. All original code used for analysis and visualization will be available from a Github repository.

## List of Supplementary Materials

Materials and Methods

Figures S1-S15

Tables S1-S19

