## Supplementary material for "Multi-omic profiling reveals widespread dysregulation of innate immunity and hematopoiesis in COVID-19": Materials and Methods; Supplementary Figures 1-15

**This PDF file includes:**

Materials and Methods
Figs. S1 to S15

**Other Supplementary Materials for this manuscript include the following:**

Tables S1 to S19

### Materials and Methods

Subjects and specimen collection

We collected blood from 64 patients enrolled in the Stanford University COVID-19 Biobanking studies from March-June 2020 after written informed consent from patients or their surrogates (Stanford IRB approvals #28205, #55650, #55689). Eligibility criteria included age ≥18 years and a positive SARS-CoV-2 nasopharyngeal swab by RT-PCR. All patients that presented to the Stanford University Emergency Department were offered enrollment regardless of admission, and patients already admitted to the wards or ICU were also eligible for enrollment. Outpatients with mild COVID-19 under the care of Stanford Health Care through the Care and Respiratory Observation of patients With novel Coronavirus (CROWN) clinic were also eligible for enrollment. The majority of admitted patients were co-enrolled in ongoing COVID-19 treatment trials at Stanford. Screening of new admissions via an electronic medical records review of all subjects was performed by the study coordinator (J.R., R.M., and H.N.D, see **Acknowledgments**) and the study principal investigators (A.J.R., S.Y., and K.C.N).

Patients were phenotyped for both peak disease severity and severity at time of sample collection according to the WHO severity score via an electronic medical records review performed by A.J.W. and D.J-M. (https://www.who.int/blueprint/priority-diseases/key-action/COVID-19_Treatment_Trial_Design_Master_Protocol_synopsis_Final_18022020.pdf). Briefly, the WHO severity score is an ordinal ranking score (0-8), where 0 indicates no evidence of infection. In this study, we classify patients with score 1-3 as "mild," corresponding to no requirement for supplemental oxygen. Score 4-5 describes "moderate" patients who are hospitalized and require non-invasive supplemental oxygen. Score 6-8 indicates "severe" infection requiring mechanical ventilation. Clinical data were obtained through the Stanford Research Repository (STARR), Stanford Medicine's approved resource for working with clinical data for research purposes extracted from the Epic database management system used by the Stanford hospitals. All fatal COVID-19 cases were confirmed by the principal investigator A.J.R. to have been primarily the result of COVID-19 and not any comorbidities.

To protect the identity of the COVID-19 subjects, ages are reported as ranges. For controls, blood was collected from eight asymptomatic adult donors as part of the Profiling Healthy Immunity study after written informed consent (Stanford IRB approval #26571). All donors were asked for consent for genetic research. Blood draws from patients occurred in concert with usual care to avoid unnecessary personal protective equipment usage. For both patients with COVID-19 and healthy controls, blood was collected into Cell Preparation Tubes (CPT) or heparin vacutainers (Becton, Dickinson and Co.) (see **Table S1**). For samples collected in CPT tubes, PBMCs were isolated by centrifugation and washed with Ca/Mg-free PBS. For samples collected in heparin tubes, PBMCs were isolated by density gradient centrifugation using Ficoll-Paque Plus medium (GE Healthcare) and washed with Ca/Mg-free PBS. For a subset of patients, whole blood was removed from CPT vacutainers prior to centrifugation and treated with ACK red blood cell lysis buffer until the cell pellet appeared visually clear, and these cells were processed for single-cell transcriptomics. Blood was processed within 6h of collection for all samples. Samples from patients with COVID-19 and healthy controls were processed side by side to avoid variation from processing. All scRNA-seq processing was performed on samples prior to cryopreservation. All CyTOF and scATAC-seq processing was performed on samples cryopreserved under the vapor phase of liquid nitrogen.

Single-cell RNA sequencing by Seq-Well

The Seq-Well platform for scRNA-seq was utilized as described previously *(15, 28, 112, 113)*. Immediately after Ficoll separation, 50,000 PBMCs were resuspended in RPMI + 10% FCS at a concentration of 75,000 cells/mL. 200 µL of this cell suspension (15,000 cells) was then loaded onto Seq-Well arrays pre-loaded with mRNA capture beads (ChemGenes). Following four washes with DPBS to remove serum, the arrays were sealed with a polycarbonate membrane (pore size of 0.01 µm) for 30 minutes at 37°C and then frozen at -80°C for no less than 24 hours and no more than 14 days to allow batching of samples processed at irregular hours. Next, arrays were thawed, cells lysed, transcripts hybridized to the mRNA capture beads, and beads recovered from the arrays and pooled for downstream processing. Immediately after bead recovery, mRNA transcripts were reverse transcribed using Maxima H-RT (Thermo Fisher EPO0753) in a template-switching-based RACE reaction, excess unhybridized bead-conjugated oligonucleotides removed with Exonuclease I (NEB M0293L), and second-strand synthesis performed with Klenow fragment (NEB M0212L) to enhance transcript recovery in the event of failed template switching *(112)*. Whole transcriptome amplification (WTA) was performed with KAPA HiFi PCR Mastermix (Kapa Biosystems KK2602) using approximately 6,000 beads per 50 µL reaction volume. Resulting libraries were then pooled in sets of 6 (approximately 36,000 beads per pool) and products purified by Agencourt AMPure XP beads (Beckman Coulter, A63881) with a 0.6x volume wash followed by a 0.8x volume wash. Quality and concentration of WTA products was determined using an Agilent Fragment Analyzer (Stanford Functional Genomics Facility), with a mean product size of >800bp and a non-existent primer peak indicating successful preparation. Library preparation was performed with a Nextera XT DNA library preparation kit (Illumina FC-131-1096) with 1 ng of pooled library using dual-index primers. Tagmented and amplified libraries were again purified by Agencourt AMPure XP beads with a 0.6x volume wash followed by a 1.0x volume wash, and quality and concentration determined by Fragment Analysis. Libraries between 400-1000bp with no primer peaks were considered successful and pooled for sequencing. Sequencing was performed on a NovaSeq 6000 instrument (Illumina; Chan Zuckerberg Biohub). The read structure was paired-end with read 1 beginning from a custom read 1 primer *(28)* containing a 12bp cell barcode and an 8 bp unique molecular identifier (UMI), and with read 2 containing 50bp of mRNA sequence.

Alignment and quality control of single-cell RNA sequencing data

Sequencing reads were aligned and count matrices assembled using STAR *(114)* and dropEst *(115)*, respectively. Briefly, the mRNA reads in read 2 demultiplexed FASTQ files were tagged with the cell barcode and UMI for the corresponding read in the read 1 FASTQ file using the dropTag function of dropEst. Next, reads were aligned with STAR using the GRCh37.p13 (hg19) human reference genome from Ensembl that included the complete genome sequences for all SARS-CoV-2 strains sequenced from California before March 24, 2020 (10 SARS-CoV-2 sequences). No SARS-CoV-2 reads were aligned from these samples using this strategy, even when the outFilterMultimapNmax behavioral option of STAR was increased from 10 (default) to 20 to accommodate for potential multiple-mapping SARS-CoV-2 reads. Count matrices were built from resulting BAM files using dropEst *(115)*. Count matrices for intron-aligned reads were also generated in order to computationally analyze cellular trajectory. Cells that had fewer than 750 UMIs or greater than 15,000 UMIs, as well as cells that contained greater than 20% of reads from mitochondrial genes or rRNA genes (*RNA18S5* or *RNA28S5*), were considered low quality and removed from further analysis. To remove putative multiplets (where more than one cell may have loaded into a given well on an array), cells that expressed more than 75 genes per 100 UMIs were also filtered out. Genes that were expressed in fewer than 10 cells were removed from the final count matrix. We removed cells with > 7.5% of reads from hemoglobin genes.

Pre-processing of single-cell RNA sequencing data

The R package Seurat *(116, 117)* was used for data scaling, transformation, clustering, dimensionality reduction, differential expression analysis, and most visualizations. Data were scaled and transformed and variable genes identified using the SCTransform() function, and linear regression performed to remove unwanted variation due to cell quality (% mitochondrial reads, % rRNA reads, % hemoglobin genes). PCA was performed using the 3,000 most highly variable genes, and the first 50 principal components (PCs) used to perform UMAP to embed the dataset into two dimensions. Next, the first 50 PCs were used to construct a shared nearest neighbor graph (SNN; FindNeighbors()) and this SNN used to cluster the dataset (FindClusters()). Although upstream quality control removed many dead or low quality cells, some clusters in the full dataset, or in cell type subsets, were identified that were defined by few canonical cell lineage markers and enriched for genes of mitochondrial or ribosomal origin, and these clusters were removed from further analysis *(118, 119)*. Manual annotation of cellular identity was performed by finding differentially expressed genes for each cluster using Seurat's implementation of the Wilcoxon rank-sum test (FindMarkers()) and comparing those markers to known cell type-specific genes from previous datasets *(120–125)*.

Automated annotation of granular cell types by Seurat v4

Seurat v4 introduced weighted nearest neighbors (WNN) analysis as a strategy to integrate multimodal single-cell sequencing data and released a large multimodal dataset of 228 cell surface markers with simultaneous whole transcriptome measurements *(30)*. By using WNN to learn the relative utility of each data modality in each cell, this reference dataset contains highly robust and granular cell type annotations. The web application provided with the release of Seurat v4 (http://azimuth.satijalab.org/) was used to map our transcriptomic dataset to the annotated multimodal reference *(30)*. Briefly, anchors between the query and reference dataset were identified using a precomputed supervised PCA on the reference dataset that maximally captures the structure of the WNN graph. Next, cell type labels from the reference dataset, as well as imputations of all measured protein markers, are transferred to each cell of the query through the previously identified anchors. Finally, the query dataset is projected onto the UMAP structure of the reference.

Calculation of transcriptomic perturbation score

To prioritize analysis of cell types of interest, we calculated a perturbation score for each cell type of each sample as previously described *(30, 31)*. This perturbation score is motivated by the observation that the statistical significance of per-gene differential expression tests is strongly influenced by the number of cells in each cluster or cell type. To overcome this, we first identified a set of genes for each cell type that showed evidence of differential expression (p value < 0.1) between healthy controls and all COVID-19 patients. From this set of genes we removed ISGs and *Ig* genes, as the former are broadly upregulated across cell types (**Fig. S13**) and the latter are cell type-, not perturbation-, specific. Next, we defined the global perturbation vector as the average log-fold change of each DEG relative to healthy controls normalized to length 1. Finally, we projected the transcriptome of each sample onto this vector and defined the perturbation score as the absolute value of the magnitude of this projection. This approach enables prioritization of cell type perturbations when comparing cell types of different abundances.

Gene module scoring analysis

The Seurat function AddModuleScore() was used to score single cells by expression of a list of genes of interest. This function calculates a module score by comparing the expression level of an individual query gene to other randomly-selected control genes expressed at similar levels to the query genes, and is therefore robust to scoring modules containing both lowly and highly expressed genes, as well as to scoring cells with different sequencing depth. Gene lists used to define each module are listed in **Table S12.** Methods to select genes from published datasets varied based on the availability, format, and modality of data. For the MDSC gene set described by Alshetaiwi, et al. *(42)*, MDSC DEGs with a log-fold change > 0.25 relative to monocytes were used. To estimate the expression of sepsis-related genes, all positively enriched genes in the MS1 module vs. MS2 module were used *(43)*. For bulk transcriptomic datasets, including LPS-stimulated PD-L1$^+$ neutrophils *(74)* and neutrophils from ARDS-complicated sepsis *(75)*, the top 97$^{th}$ percentile of differentially expressed genes relative to control samples/patients were used for scoring.

Transcription factor activity prediction analysis

The iRegulon plugin available through Cytoscape was used to predict transcription factors that may contribute to the observed transcriptomic changes. Gene lists supplied to iRegulon are available in **Table S14.** In brief, iRegulon calculates the likelihood of transcription factor activity first by compiling for each transcription factor motif a ranked list of genes containing that motif near the TSS, and then determining the ability of each transcription factor motif to recover the genes supplied as input. Transcription factors with a normalized enrichment score of > 4 and > 20 predicted targets, along with their corresponding predicted regulomes, are plotted for visualization.

Analysis of developing neutrophil trajectories

PHATE *(77)* was performed on scaled and transformed expression values for the 3,000 most highly variable genes to embed the subsetted dataset into two or three dimensions. RNA velocity analysis was performed with the package scVelo to visualize RNA velocity field vectors and streams as well as to calculate the latent time of each developing neutrophil. Plots of expression of individual genes along inferred latent time are scaled at the 5$^{th}$ and 95$^{th}$ percentiles.

Projection of transcriptomic dataset onto published blood and bone marrow hematopoiesis dataset

Cells annotated as HSPCs by Seurat v4 were projected into a publicly available hematopoiesis dataset of CD34$^+$-enriched BMMCs *(90)* using an anchor-based integration strategy. Briefly, expression values from each dataset were normalized and variable features identified using SCTransform without covariate regression. Next, anchors were identified between the two datasets, the datasets integrated using these anchors, and PCA and UMAP performed as described above on the integrated gene expression matrix. To determine the identities of the projected cells, TransferData() was used to transfer cell type labels from the cells from the published dataset in the integrated object to the HSPCs from the COVID-19 dataset.

Mortality prediction using developing neutrophil DEGs

To test whether the gene signature of developing neutrophils could be used to predict COVID-19 mortality, we first developed a five-gene signature of developing neutrophils by identifying the most differentially-enriched genes in developing neutrophils in our transcriptomic dataset relative to all other cells. Next, we downloaded normalized transcript counts from a publicly-available whole blood bulk transcriptomic dataset published by Overmyer, et al. *(91)*. We then scored each COVID-19 sample in this dataset by the expression of the five developing neutrophil-enriched genes (*DEFA1B*, *DEFA3*, *LTF*, *DEFA1*, and *S100A8*) using AddModuleScore(). Finally, we used these gene signature scores as a predictor variable and 28-day mortality as reported by metadata from Overmyer, et al. as the response variable to construct an ROC curve to quantify and visualize the sensitivity and specificity of the prediction.

NK Cell Isolation

After thawing of PBMC, 0.5e6 live PBMC were set aside from each sample for staining with a broad lineage panel that includes NK cell ligands (ligand panel) and the remaining cells were used for NK cell isolation. NK cells were isolated from whole PBMC via Miltenyi Human NK Cell Isolation Kits, which utilize negative selection, per the manufacturer's instructions.

Mass Cytometry

All antibodies not purchased directly from Fluidigm were conjugated using MaxPar X8 conjugation kits (Fluidigm). To ensure staining consistency, all antibodies except those noted were pre-combined into staining cocktails and either lyophilized (NK surface and ICS panels) or frozen at -80C (Ligand panel) for long-term storage. Samples were barcoded with a four-choose-two scheme, utilizing Palladium isotopes (Pd102, Pd104, Pd106, Pd108) conjugated to anti-CD45 antibodies as previously described *(126)*.

Following isolation of NK cells, both the whole PBMC and the isolated NK cells from each sample were stained with Cisplatin for one minute in order to allow for viability determination and subsequently quenched with FBS. Samples were then stained with Palladium-CD45 barcodes as previously described (JoVE paper). After barcode staining at 4 C, samples were washed thoroughly with CyFACS buffer (PBS, 0.1% BSA, 2mM EDTA, 0.05% sodium azide) and combined into sets of barcodes, hereafter referred to as "barcoded samples". Barcoded samples were subsequently washed and stained with relevant surface panels (**Fig. S15A-B**) for 30 minutes at 4 C. Following surface stain, the barcoded samples were washed with CyFACS buffer and fixed in 2% Paraformaldehyde for 20 minutes at room temperature. Barcoded samples were subsequently permeabilized with 1x BD Perm II (BD Biosciences, Franklin Lakes, NJ, USA) and NK cell samples were stained with the lyophilized ICS panel for 45 minutes at 4 C (whole PBMC were resuspended in 1x BD Perm II and left at 4 C for the same duration without any stains). All of the barcoded samples were then resuspended in iridium interchelator (DVS Sciences) in 2% PFA until collection (within 3 days of staining).

Data was collected on a Helios mass cytometer. Prior to collection, samples were washed with CyFACS buffer and Milli-Q water before being resuspended in 1x EQ beads (Fluidigm) for collection.

Preprocessing and Data Analysis of Mass Cytometry Data

FCS files were normalized and debarcoded using the Premessa package as previously described *(127, 128)*. FlowJo v10.7.1 was used to manually gate out EQ beads, dead cells, doublets, and debris from both whole PBMC and NK cell samples. Additional lineage markers were used to exclude contaminating non-NK cells from the samples consisting of bead-purified NK cells. Live, intact, singlet PBMC were exported from whole PBMC samples as FCS files while live, intact, singlet NK cells were exported from NK cell samples using the gating schemes described in **Fig. S15C-D**. All subsequent preprocessing and downstream analysis of data was performed using the open-source software R (**Fig. S15e**). NK cell fcs files were normalized to account for batch effects using the *CytoNorm* package *(129)*; this normalization was not performed on the whole PBMC files as no such batch effects were observed. Parameter names were altered in whole PBMC fcs files using the *Premessa* panel editor tool to ensure consistency across all samples. Seurat objects were created from FCS files using the *Seurat* and *FlowCore* packages. Unsupervised clustering was performed on the whole PBMC Seurat object via the PARC algorithm *(130)*, using all proteomic parameters listed in **Fig. S15A** with the exception of HLA-Bw4 and HLA-Bw6. These parameters were excluded as their expression is mutually exclusive and can therefore drive overclustering. Clustering was performed on whole PBMC with a resolution of 0.19 and on subsetted monocyte clusters with a resolution of 0.21. Clustering resolutions were determined using cluster tree plots (**Fig. S2G, Fig. S10B**). Clustering and UMAP embeddings were performed on arcsinh-transformed data (cofactor = 5). All boxplots depicting CyTOF data represent transformed per-sample mean signal.

Single-cell ATAC (assay for transposase-accessible chromatin) sequencing

The single-cell ATAC sequencing was performed by using the Chromium Next GEM Single Cell ATAC Reagent Kits v1.1 (10x Genomics, PN-1000175) and following the Demonstrated Protocol (Nuclei Isolation for Single Cell ATAC Sequencing) provided by the 10x Genomics company. Briefly, cryopreserved PBMCs were thawed and 50,000-500,000 cells were aliquoted, washed with PBS and then lysed with 100 µl lysis buffer for 4 min. The lysed nuclei were centrifuged after washing with 1 ml washing buffer, and resuspended in Diluted Nuclei buffer. The nuclei concentration was then determined with TC20™ Automated Cell Counter (Bio-rad, #1450102), and approximately 6,000 nuclei were used for Tn5 transposition, single nuclei barcoding and library preparation following the instructions in the kit. The final DNA libraries were sequenced with NovaSeq 6000 system (Illumina; Chan Zuckerberg Biohub), leading to around 200 million reads per sample.

scATAC-seq preprocessing and cell quality filtering

Demultiplexed sequencing reads were aligned to the GRCh37 (hg19) human reference genome using cellranger-atac software (10x Genomics, v1.2), or cellranger-arc (10x Genomics, v1.0) for the multi-omic reference dataset. Resulting fragment files were processed using ArchR *(131)*, and filtered based off of TSS enrichment and the log10 fragments per cell. Cell quality cutoff values were set on a per-sample basis to account for variable sequencing depths and sample qualities. TSS cutoff ranged from 5-10 , and log10 fragments cutoff ranged from 3.2-4.1 such that the large number of non-cell-containing droplets were excluded from further consideration (**Table S17**). Cross-celltype doublets were computationally identified using ArchR's addDoubletScores function, and filtered based on a maximum doublet enrichment of 2 (i.e. regions of the manifold that are 2-fold enriched for simulated doublets).

scATAC-seq reference-based cell type annotation

A public multi-omic dataset from 10x genomics on healthy PBMCs was used as an intermediate for cell type calls (https://support.10xgenomics.com/single-cell-multiome-atac-gex/datasets/1.0.0/pbmc_granulocyte_sorted_10k), as it could be readily integrated with both scRNA-seq and scATAC-seq datasets. The multi-omic dataset was re-aligned to hg19 using cellranger-arc v1.0 (10x Genomics), and the resulting RNA counts matrix was filtered for cells with between 1,000 and 15,000 reads, and under 20% mitochondrial genes. Next, cell types were annotated using the Azimuth tool from Seurat v4 in the same manner as the scRNA dataset *(30)*. Macs2 was used to call peaks for each the Azimuth-annotated cell types via ArchR's addReproduciblePeakSet, a peak counts matrix was calculated using addPeakMatrix, then the peak matrix was dimensionality reduced using the addIterativeLSI function from ArchR with iterations=3, varFeatures=50000, and sampleCellsPre=50000. This established an ATAC-based dimensionality reduction coupled to RNA-based automated cell type annotations.

Each scATAC sample from our dataset was then projected into the linear dimensionality reduction of the multi-omic dataset. Then Seurat's internal FindAnchors function and TransferData functions *(116)* were used to transfer cell type annotations from the multi-omic dataset to each scATAC dataset. Briefly, anchor pairs between the datasets are identified based on mutual nearest neighbors, then cell type calls are transferred for each cell using a distance-weighted sum of nearby anchors. The anchor filtering step was omitted (k.filter=NA) as suggested in the scATAC-seq data integration vignette from Signac *(132)*. This avoids over-filtering of anchors due to the extremely sparse nature of the underlying ATAC-seq data.

scATAC Batch Correction and Dimensionality Reduction

Cells passing quality and doublet filters from each sample were combined into a linear dimensionality reduction using ArchR's addIterativeLSI function. This dimensionality reduction was then corrected for batch effect based on processing date using the Harmony method *(133)*, via ArchR's addHarmony function. The cells were then clustered based on the batch-corrected dimensions using ArchR's addClusters function -- briefly, this uses a modularity-based clustering of the shared-nearest-neighbor graph as implemented in Seurat. Three doublet clusters were manually identified from these clusters, based on containing a mixture of many cell types and elevated doublet enrichment scores (although still below the doublet cutoff threshold of 2). These doublet clusters were removed from further consideration in all downstream analyses.

scATAC Transcription Factor Activity Analysis

For transcription factor activity analysis in CD14 monocytes, peaks were called on CD14 monocyte cells using the addReproduciblePeakSet from ArchR. Then, peaks containing NF-κB2 motifs were identified using the addPeakAnnotations function, which relies on the chromVARmotifs package's human_pwms_v2 motif set curated from the cisBP database. Motif deviation z-scores for individual cells were calculated using ArchR's addDeviationsMatrix function. Briefly, this method compares accessibility across all peaks containing a transcription-factor motif to accessibility across a background set of peaks matched for GC content and overall accessibility. This measures global changes in accessibility associated with transcription factor activity while controlling for technical confounders.

Hi-C data

Hi-C data for monocytes was taken from Phanstiel, et al. *(51)*, downloaded using juicer tools v1.22 *(134)*.

Data visualization

Wrappers provided by Seurat were used to generate UMAP projections and Dot Plots. ComplexHeatmap was used to generate all heatmaps, and plotly used to visualize three dimensional PHATE projections. scVelo was used to visualize RNA velocity streams. Custom ggplot functions (see **Data and materials availability**) were used to generate all other plots. For all boxplots, features include: minimum whisker, 25th percentile − 1.5 × inter-quartile range (IQR) or the lowest value within; minimum box, 25th percentile; center, median; maximum box, 75th percentile; maximum whisker, 75th percentile + 1.5 × IQR or greatest value within.

### Supplementary Figures

A

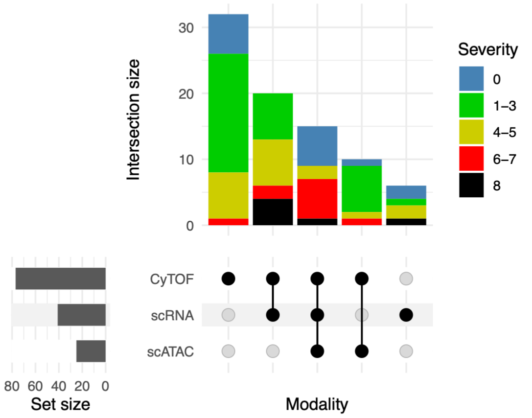


B

| Demographics | 0 | 1-3 | 4-5 | 6-7 | 8 |
|---|---|---|---|---|---|
| Number | 14 | 33 | 19 | 10 | 6 |
| Age [Median (Q1-Q3)] | 46.5 (35-49) | 36 (30-55) | 58 (41-67) | 40 (36-62) | 61 (46-74) |
| Male [%] | 50.0 | 39.4 | 47.4 | 60.0 | 83.3 |
| **Ethnicity %** | **0** | **1-3** | **4-5** | **6-7** | **8** |
| Caucasian | 75.0 | 24.2 | 21.1 | 10.0 | 16.7 |
| Black | 0.0 | 6.1 | 5.3 | 10.0 | 0.0 |
| Asian | 25.0 | 6.1 | 5.3 | 30.0 | 0.0 |
| Latinx | 0.0 | 54.5 | 63.2 | 50.0 | 66.7 |
| Other/unknown | 0.0 | 9.1 | 5.3 | 0.0 | 16.7 |
| **Clinical course** | **0** | **1-3** | **4-5** | **6-7** | **8** |
| BMI [Median (Q1-Q3)] | -- | 26.1 (24.3-30.1) | 33.5 (29.3-36.2) | 35.2 (31.9-39.9) | 29.2 (26.2-34.0) |
| Dialysis [%] | 0.0 | 0.0 | 0.0 | 10.0 | 33.3 |
| Days in hospital [Median (Q1-Q3)] | -- | 0.3 (0.2-0.6) | 8.4 (4.5-11.8) | 29.3 (19.8-48.3) | 35.5 (18.7-39.5) |
| Days post-positive swab [Median (Q1-Q3)] | -- | 3.0 (1.0-34.0) | 2.0 (1.0-7.0) | 2.0 (1.3-2.8) | 1.0 (0.3-2.5) |
| Days before death [Median (Q1-Q3)] | -- | -- | -- | -- | 34.5 (18.8-38.3) |

C

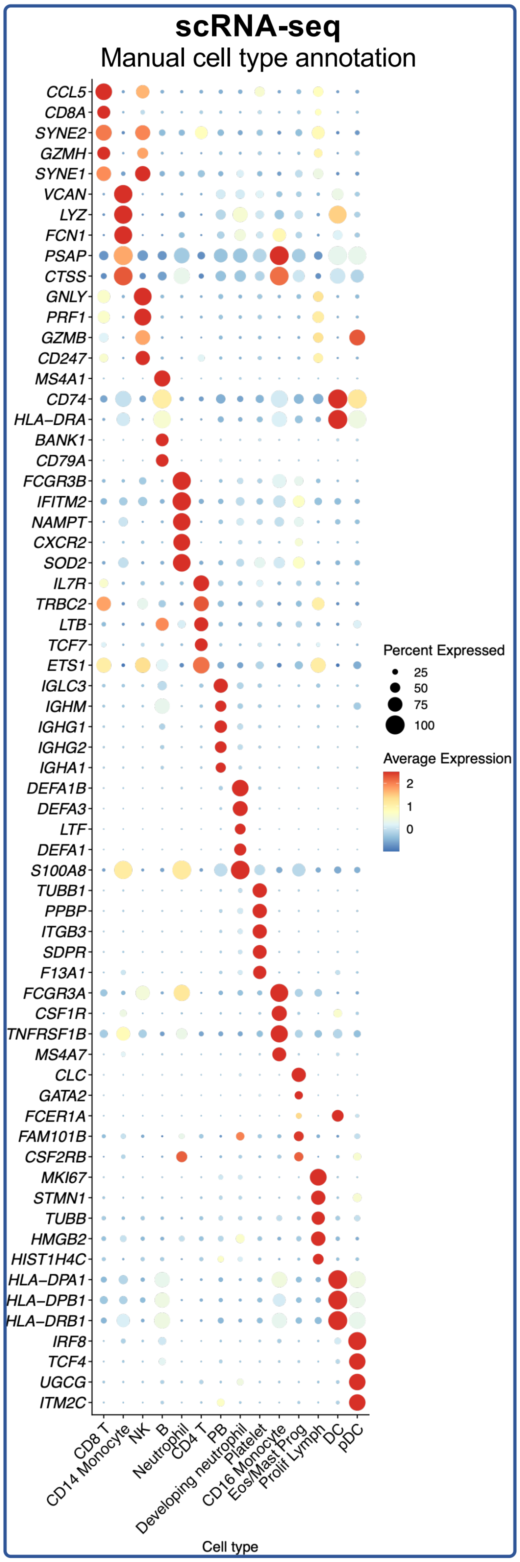


**Fig. S1. Overview of COVID-19 patient cohort and manual annotation of scRNA-seq dataset. A)** Upset plot depicting overlap of patient samples profiled between the three modalities, colored by peak disease severity score. **B)** Table summarizing key demographic and clinical features of profiled patients. **C)** Dot plot depicting percent and average expression of the top five DEGs for each manually-annotated cell type (see **Methods, Table S6**).

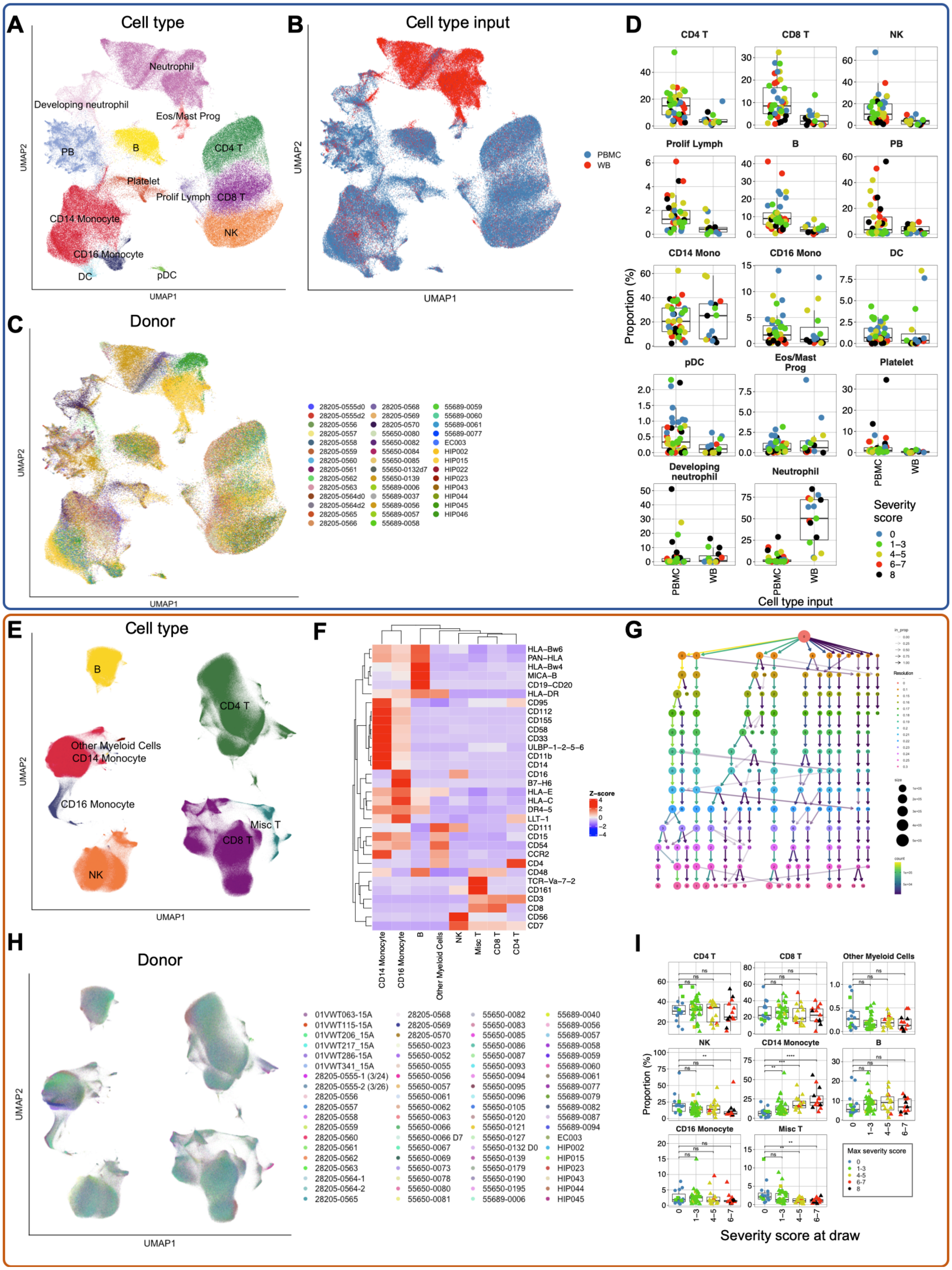
A
Cell type
Neutrophil
Developing neutrophil
Eos/Mast Prog
PB
B
CD4 T
Platelet
Prolif Lymph
CD8 T
CD14 Monocyte
NK
CD16 Monocyte
DC
pDC
UMAP2
UMAP1
B
Cell type input
PBMC
WB
C
Donor
28205-0555d0
28205-0555d2
28205-0556
28205-0557
28205-0558
28205-0559
28205-0560
28205-0561
28205-0562
28205-0563
28205-0564d0
28205-0564d2
28205-0565
28205-0566
28205-0568
28205-0569
28205-0570
55650-0080
55650-0082
55650-0084
55650-0085
55650-0132d7
55650-0139
55689-0006
55689-0037
55689-0056
55689-0057
55689-0058
55689-0059
55689-0060
55689-0061
55689-0077
EC003
HIP002
HIP015
HIP022
HIP023
HIP043
HIP044
HIP045
HIP046
D
CD4 T
CD8 T
NK
Prolif Lymph
B
PB
CD14 Mono
CD16 Mono
DC
pDC
Eos/Mast Prog
Platelet
Developing neutrophil
Neutrophil
Proportion (%)
PBMC
WB
Cell type input
Severity score
0
1-3
4-5
6-7
8
E
Cell type
B
CD4 T
Other Myeloid Cells
CD14 Monocyte
CD16 Monocyte
Misc T
CD8 T
NK
UMAP2
UMAP1
F
HLA-Bw6
PAN-HLA
HLA-Bw4
MICA-B
CD19-CD20
HLA-DR
CD95
CD112
CD155
CD58
CD33
ULBP-1-2-5-6
CD11b
CD14
CD16
B7-H6
HLA-E
HLA-C
DR4-5
LLT-1
CD111
CD15
CD54
CCR2
CD4
CD48
TCR-Va-7-2
CD161
CD3
CD8
CD56
CD7
Z-score
CD14 Monocyte
CD16 Monocyte
B
Other Myeloid Cells
NK
Misc T
CD8 T
CD4 T
G
H
Donor
01VWT063-15A
01VWT115-15A
01VWT206_15A
01VWT217_15A
01VWT286-15A
01VWT341_15A
28205-0555-1 (3/24)
28205-0555-2 (3/26)
28205-0556
28205-0557
28205-0558
28205-0559
28205-0560
28205-0561
28205-0562
28205-0563
28205-0564-1
28205-0564-2
28205-0565
28205-0568
28205-0569
28205-0570
55650-0023
55650-0052
55650-0055
55650-0056
55650-0057
55650-0061
55650-0062
55650-0063
55650-0066
55650-0066 D7
55650-0067
55650-0069
55650-0073
55650-0078
55650-0080
55650-0081
55650-0082
55650-0083
55650-0085
55650-0086
55650-0087
55650-0093
55650-0094
55650-0095
55650-0096
55650-0105
55650-0120
55650-0121
55650-0127
55650-0132 D0
55650-0139
55650-0179
55650-0190
55650-0195
55689-0006
55689-0040
55689-0056
55689-0057
55689-0058
55689-0059
55689-0060
55689-0061
55689-0077
55689-0079
55689-0082
55689-0087
55689-0094
EC003
HIP002
HIP015
HIP023
HIP043
HIP044
HIP045
I
CD4 T
CD8 T
Other Myeloid Cells
NK
CD14 Monocyte
B
CD16 Monocyte
Misc T
Proportion (%)
Max severity score
Severity score at draw

**Fig. S2. Overview of cell types and proportions present in the scRNA-seq and CyTOF dataset. A-C**) UMAP embeddings of complete scRNA-seq dataset, colored by cell subset (**A**), cell type input (**B**; either PBMCs or ACK-lysed whole blood) or donor (**C**) **D**) Box plots showing the proportion of each manually-annotated cell subset within the scRNA-seq dataset, grouped by cell type input. Each point represents the proportion of the cell type of interest in one donor; points are colored by maximum severity score. **E**) UMAP embedding of the complete PBMC CyTOF dataset, colored by cell subset. **F**) Heatmap showing the z-score normalized average expression of each marker in the PBMC CyTOF panel across all cell subsets detected in that dataset. **G**) Cluster tree diagram illustrating the different clusters of cells that resulted from performing PARC at different resolution values. A resolution of 0.19 was chosen for unsupervised analysis based on the relative cluster stability achieved at this point (see **Methods**). **H**) UMAP embedding of our whole PBMC CyTOF dataset, colored by donor. **I**) Box plots depicting the proportion of each cell subset in the PBMC CyTOF dataset, grouped by severity score at the time of sample collection and colored by maximum severity score. *, $p < 0.05$; **, $p < 0.01$; ***, $p < 0.001$; ****, $p < 0.0001$, n.s., not significant at $p = 0.05$ by two-sided Wilcoxon rank-sum test with Bonferroni's correction for multiple hypothesis testing.

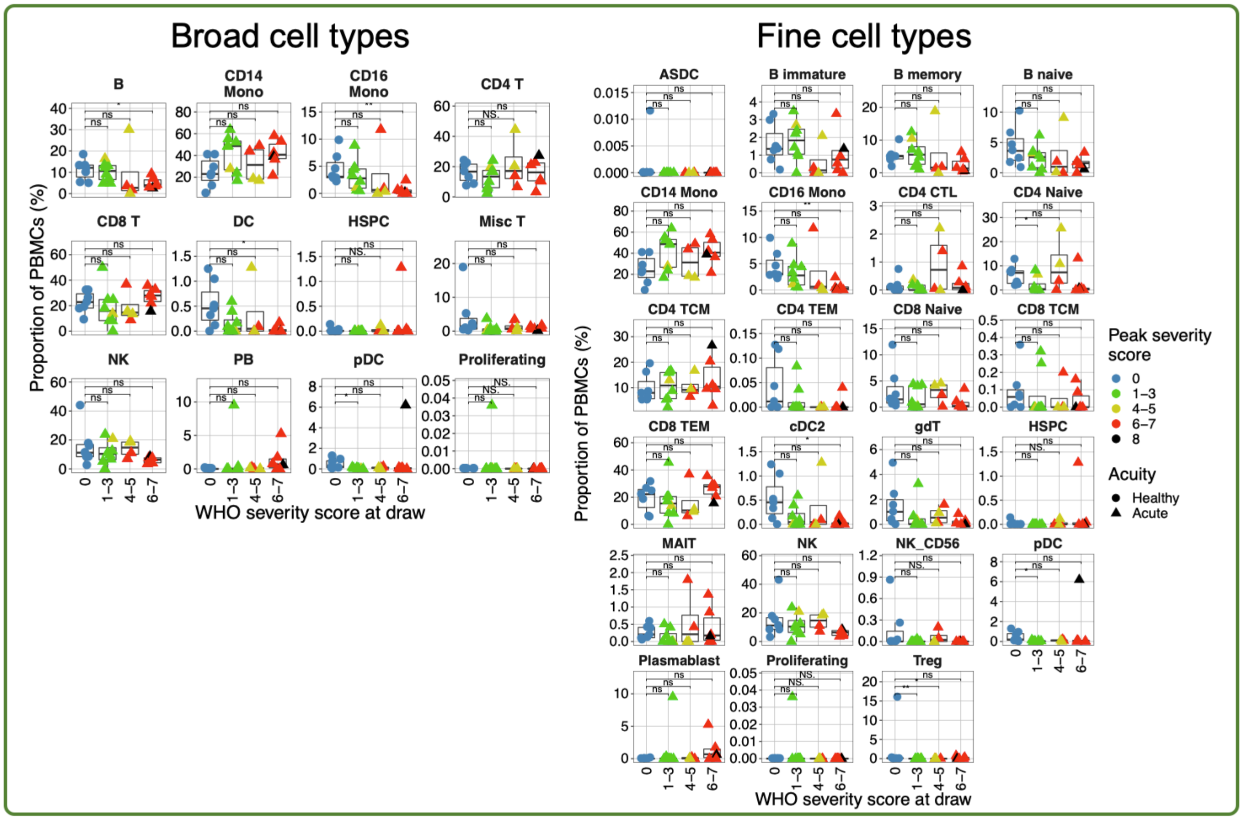


**Fig. S3**. **Cell type proportions in scATAC-seq dataset**. Box plots depicting the proportion of broad cell types (left) or fine cellular subtypes transferred from the Seurat v4 multimodal reference (right; see **Methods**). Points are colored by the peak disease severity score, shaped according to disease acuity, and grouped by the disease severity score at time of sample collection. *, p < 0.05; **, p < 0.01; ***, p < 0.001; ****, p < 0.0001, n.s., not significant at p = 0.05 by two-sided Wilcoxon rank-sum test with Bonferroni's correction for multiple hypothesis testing.

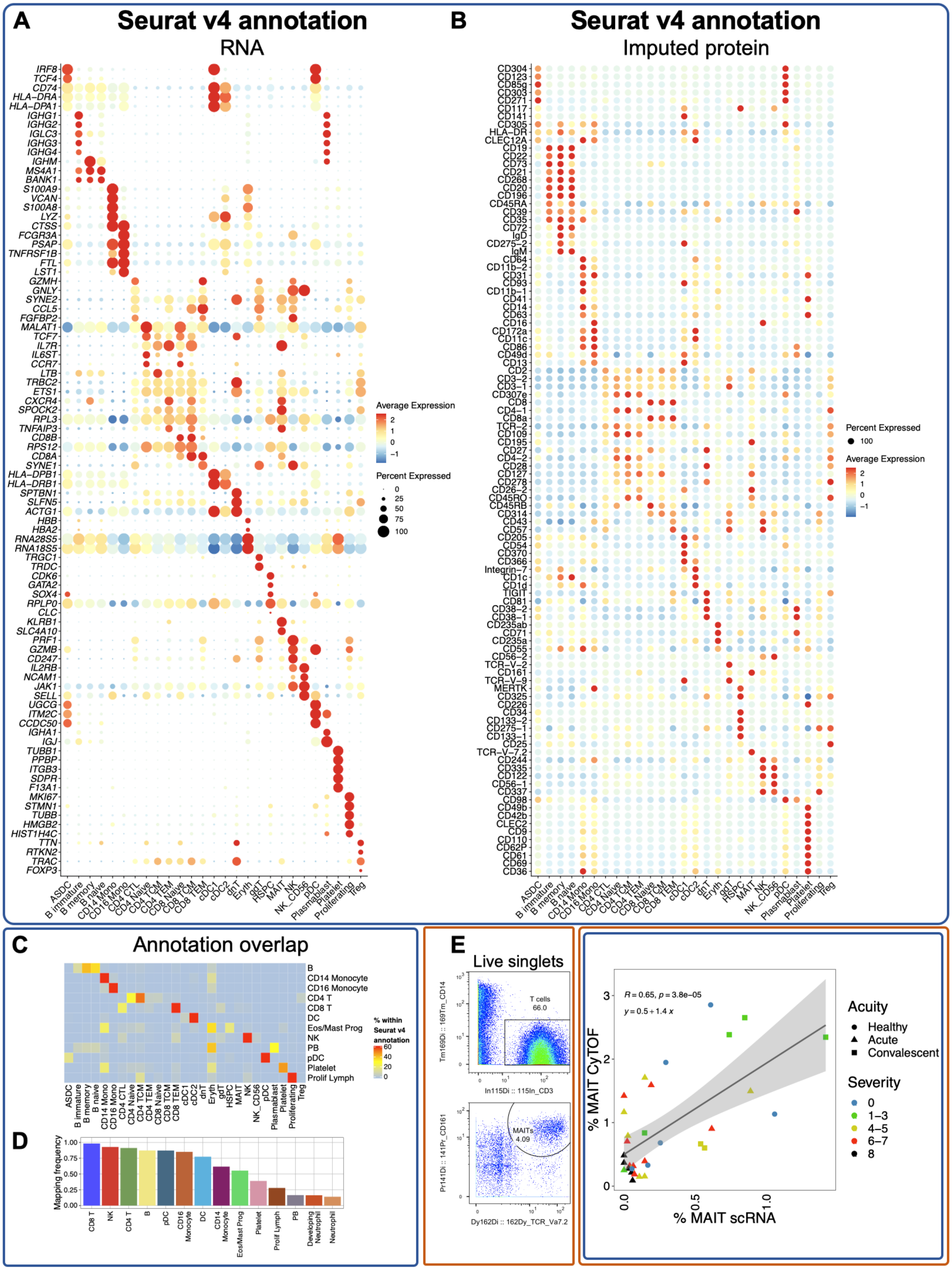

A
Seurat v4 annotation
RNA
IRF8
TCF4
CD74
HLA−DRA
HLA−DPA1
IGHG1
IGHG2
IGLC3
IGHG3
IGHG4
IGHM
MS4A1
BANK1
S100A9
VCAN
S100A8
LYZ
CTSS
FCGR3A
PSAP
TNFRSF1B
FTL
LST1
GZMH
GNLY
SYNE2
CCL5
FGFBP2
MALAT1
TCF7
IL7R
IL6ST
CCR7
LTB
TRBC2
ETS1
CXCR4
SPOCK2
RPL3
TNFAIP3
CD8B
RPS12
CD8A
SYNE1
HLA−DPB1
HLA−DRB1
SPTBN1
SLFN5
ACTG1
HBB
HBA2
RNA28S5
RNA18S5
TRGC1
TRDC
CDK6
GATA2
SOX4
RPLP0
CLC
KLRB1
SLC4A10
PRF1
GZMB
CD247
IL2RB
NCAM1
JAK1
SELL
UGCG
ITM2C
CCDC50
IGHA1
IGJ
TUBB1
PPBP
ITGB3
SDPR
F13A1
MKI67
STMN1
TUBB
HMGB2
HIST1H4C
TTN
RTKN2
TRAC
FOXP3
ASDC
B immature
B memory
B naive
CD14 Mono
CD16 Mono
CD4 CTL
CD4 Naive
CD4 TCM
CD4 TEM
CD8 Naive
CD8 TCM
CD8 TEM
cDC1
cDC2
dnT
Eryth
gdT
HSPC
MAIT
NK
NK_CD56
pDC
Plasmablast
Platelet
Proliferating
Treg
Average Expression
2
1
0
−1
Percent Expressed
0
25
50
75
100
B
Seurat v4 annotation
Imputed protein
CD304
CD123
CD85g
CD303
CD271
CD117
CD141
CD305
HLA−DR
CLEC12A
CD19
CD22
CD73
CD21
CD268
CD20
CD196
CD45RA
CD39
CD35
CD72
IgD
CD275−2
IgM
CD64
CD11b−2
CD31
CD93
CD11b−1
CD41
CD14
CD63
CD16
CD172a
CD11c
CD86
CD49d
CD13
CD2
CD3−2
CD3−1
CD307e
CD8
CD4−1
CD8a
TCR−2
CD109
CD195
CD27
CD4−2
CD28
CD127
CD278
CD26−2
CD45RO
CD45RB
CD314
CD43
CD57
CD205
CD54
CD370
CD366
Integrin−7
CD1c
CD1d
TIGIT
CD81
CD38−2
CD38−1
CD235ab
CD71
CD235a
CD55
CD56−2
TCR−V−2
CD161
TCR−V−9
MERTK
CD325
CD226
CD34
CD133−2
CD275−1
CD133−1
CD25
TCR−V−7.2
CD244
CD335
CD122
CD56−1
CD337
CD98
CD49b
CD42b
CLEC2
CD9
CD110
CD62P
CD61
CD69
CD36
Percent Expressed
100
Average Expression
2
1
0
−1
C
Annotation overlap
B
CD14 Monocyte
CD16 Monocyte
CD4 T
CD8 T
DC
Eos/Mast Prog
NK
PB
pDC
Platelet
Prolif Lymph
% within Seurat v4 annotation
60
40
20
0
D
Mapping frequency
1.00
0.75
0.50
0.25
0.00
CD8 T
NK
CD4 T
B
pDC
CD16 Monocyte
DC
CD14 Monocyte
Eos/Mast Prog
Platelet
Prolif Lymph
PB
Developing Neutrophil
Neutrophil
E
Live singlets
T cells
66.0
Tm169Di :: 169Tm_CD14
In115Di :: 115In_CD3
MAITs
4.09
Pr141Di :: 141Pr_CD161
Dy162Di :: 162Dy_TCR_Va7.2
R = 0.65, p = 3.8e−05
y = 0.5 + 1.4 x
% MAIT CyTOF
% MAIT scRNA
Acuity
Healthy
Acute
Convalescent
Severity
0
1−3
4−5
6−7
8

**Fig. S4. Seurat v4 robustly annotates and predicts protein expression of biologically-relevant cell types**. **A-B)** Dot plot depicting average and percent expression of the top five DEGs (**A**) or top five differentially-abundant imputed protein markers (**B**) for each Seurat v4-annotated cell type. **C)** Heatmap showing overlap in Seurat v4 annotation calls (*x* axis) and manual cell type annotations (*y* axis), colored by the percentage of a manual cell annotation within a Seurat v4 annotation (ie. each row sums to 100%). **D)** Bar plot showing mapping frequency of each manually assigned cell type annotation by Seurat v4. Neutrophils and developing neutrophils are the least frequently assigned cell types as they are not present in the reference dataset *(30)*. **E)** Manual gating scheme for MAIT cells in CyTOF dataset, beginning with live singlets gated according to the scheme in **Fig. S15. F)** Scatter plot depicting concordance with proportions of MAITs predicted by Seurat v4 in the scRNA-seq dataset (*x* axis) and proportions of MAITs manually gated in the CyTOF dataset (*y* axis). Pearson's r and exact two-sided p values are shown for the correlation, as well as the equation for a line of best fit.

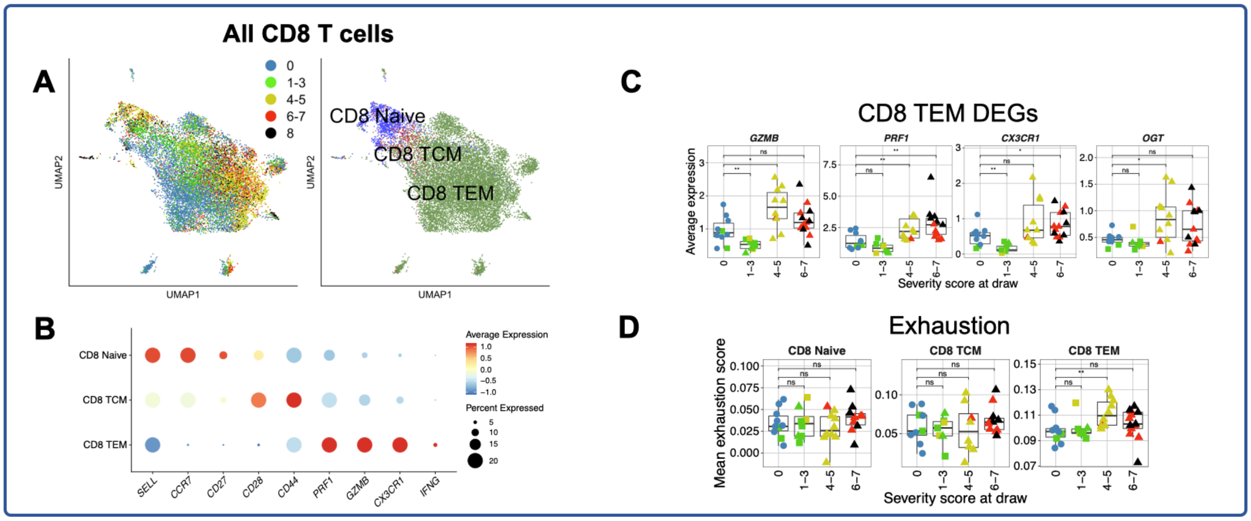


**Fig. S5. Disease severity-associated perturbations in CD8 $T_{EM}$ cells. A)** UMAP projection of all CD8 T cells colored by peak disease severity score (left) and Seurat v4-annotated cell type (right). **B)** Dot plot depicting percent and average expression of canonical CD8 subset-defining genes (see **Methods, Fig. S4a**). **C)** Box plots depicting average expression of selected DEGs (see **Table S18** for complete list) by CD8 $T_{EM}$ cells in each sample. **D)** Box plots showing average module scores for T cell exhaustion (as reported in *(135)*, see **Table S12**) in each annotated CD8 T cell subset. For all box plots: points are colored by the peak disease severity score, shaped according to disease acuity, and grouped by the disease severity score at time of sample collection. *, $p < 0.05$; **, $p < 0.01$; ***, $p < 0.001$; ****, $p < 0.0001$, n.s., not significant at $p = 0.05$ by two-sided Wilcoxon rank-sum test with Bonferroni's correction for multiple hypothesis testing.

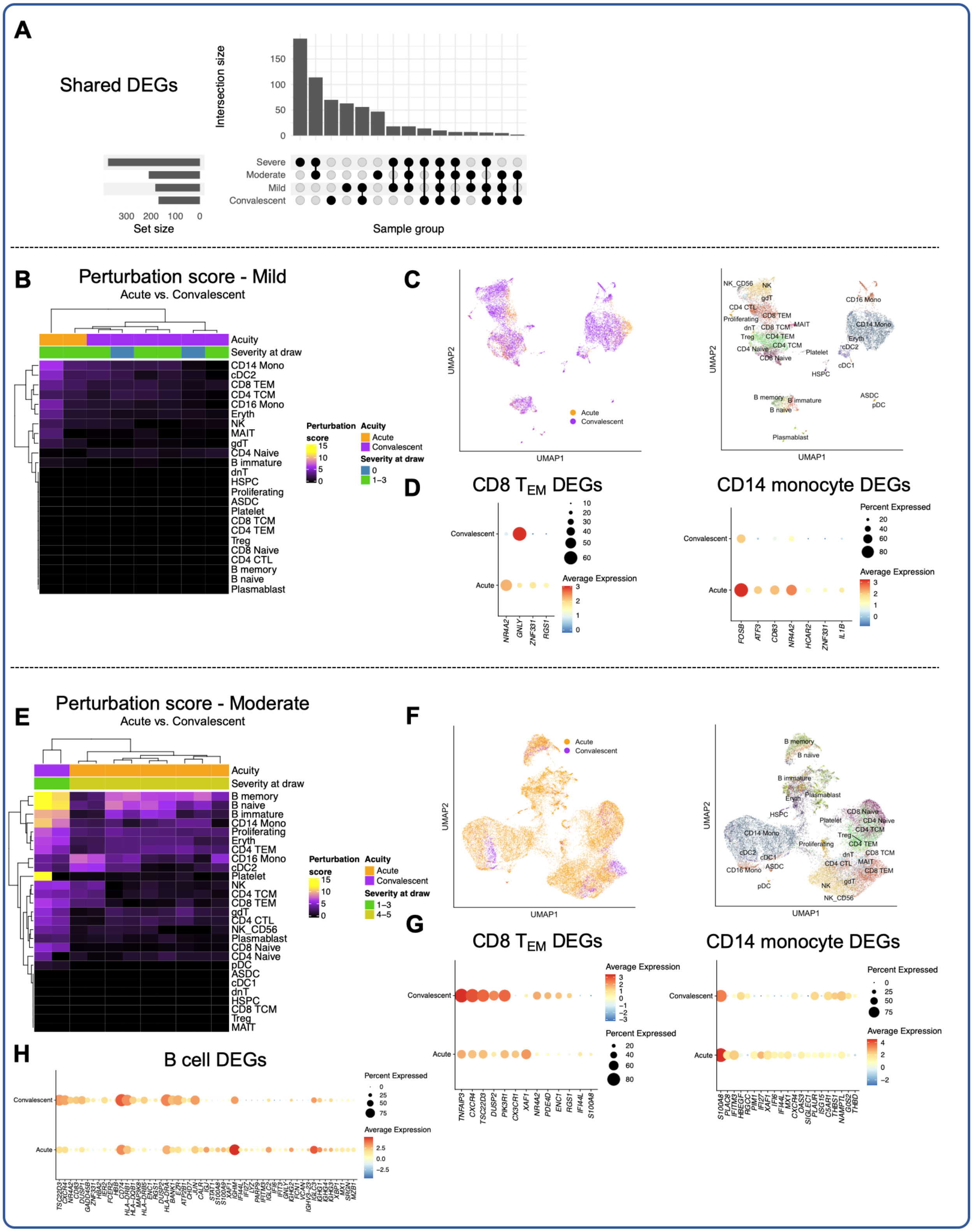


**Fig. S6. Impact of disease acuity on transcriptomic phenotype of mild and moderate COVID-19 patients. A)** Upset plot depicting overlap of DEGs between acute mild, acute moderate, acute severe, and convalescent samples when each is compared to healthy controls.

DEGs testing is performed on PBMCs and filtered for adjusted p value < 1e-4. **B,E)** Heatmaps of cellular perturbation score, as described by Papalexi, et al. *(31)*, per mild (**B**) or moderate (**E**) COVID-19 sample in each Seurat v4-labeled cell type. DEGs between mild vs. convalescent samples in each severity group are used as input for each perturbation score (see **Methods**). **C,F)** UMAP projections of all cells from mild (**C**) or moderate (**F**) COVID-19 patients colored by disease acuity (left) and Seurat v4-annotated cell type (right). **D,G)** Dot plots depicting percent and unscaled average expression for all DEGs with |log(fold-change| > 1 in CD8 $T_{EM}$ cells (left) and CD14 monocytes (right) of mild (**D**) or moderate (**G**) COVID-19 patients. **H)** Dot plot depicting percent and unscaled average expression for all DEGs with |log(fold-change| > 1 in B cells of moderate COVID-19 patients.

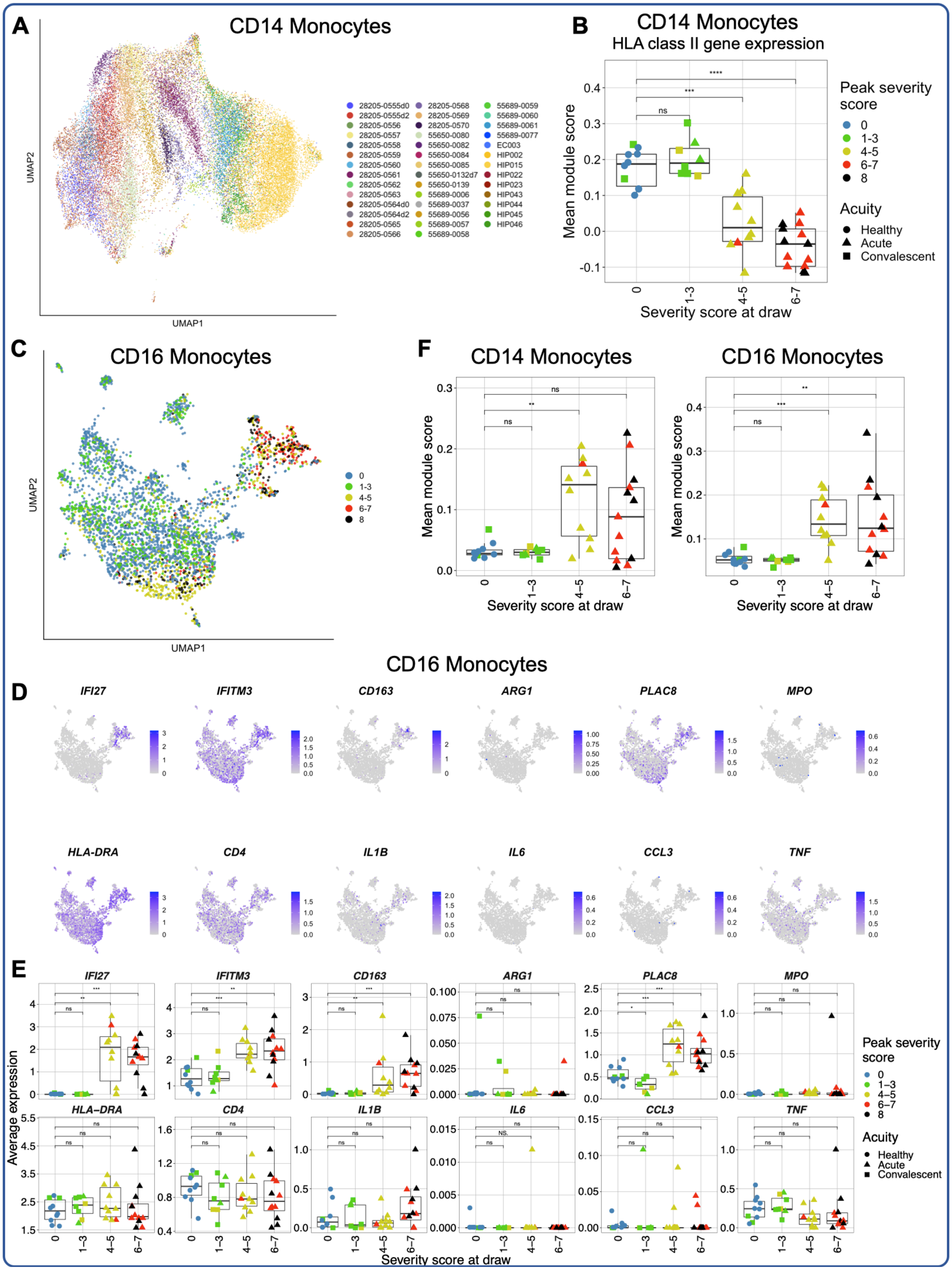
A
CD14 Monocytes
UMAP1
UMAP2
B
CD14 Monocytes
HLA class II gene expression
Mean module score
Severity score at draw
Peak severity score
0
1-3
4-5
6-7
8
Acuity
Healthy
Acute
Convalescent
C
CD16 Monocytes
F
CD14 Monocytes
CD16 Monocytes
CD16 Monocytes
D
IFI27
IFITM3
CD163
ARG1
PLAC8
MPO
HLA-DRA
CD4
IL1B
IL6
CCL3
TNF
E
Average expression
Severity score at draw

**Fig. S7. Additional analysis of CD14 and CD16 monocyte transcriptomic phenotype in COVID-19. A)** UMAP embedding of all $CD14^+$ monocytes in the scRNA-seq dataset, colored by donor. **B)** Box plot showing median module expression of HLA Class II genes in $CD14^+$ monocytes (see **Methods, Table S12**). **C-D)** UMAP projection of all $CD16^+$ monocytes in the scRNA-seq dataset, colored by maximum WHO severity score **(C)** or by expression of $CD14^+$ monocyte DEGs depicted in **Fig. 3** (*IF127, IFITM3, CD163, ARG1, PLAC8, MPO, HLA-DRA, CD4, IL1B, IL6, CCL3,* and *TNF*) **(D)**. **E)** Box plots quantifying the average expression per donor of genes visualized in **(D)**. **F)** Average ISG module score per sample in $CD14^+$ (left) and $CD16^+$ (right) monocytes. For all box plots: points are colored by the peak disease severity score, shaped according to disease acuity, and grouped by the disease severity score at time of sample collection. *, $p < 0.05$; **, $p < 0.01$; ***, $p < 0.001$; ****, $p < 0.0001$, n.s., not significant at $p = 0.05$ by two-sided Wilcoxon rank-sum test with Bonferroni's correction for multiple hypothesis testing.

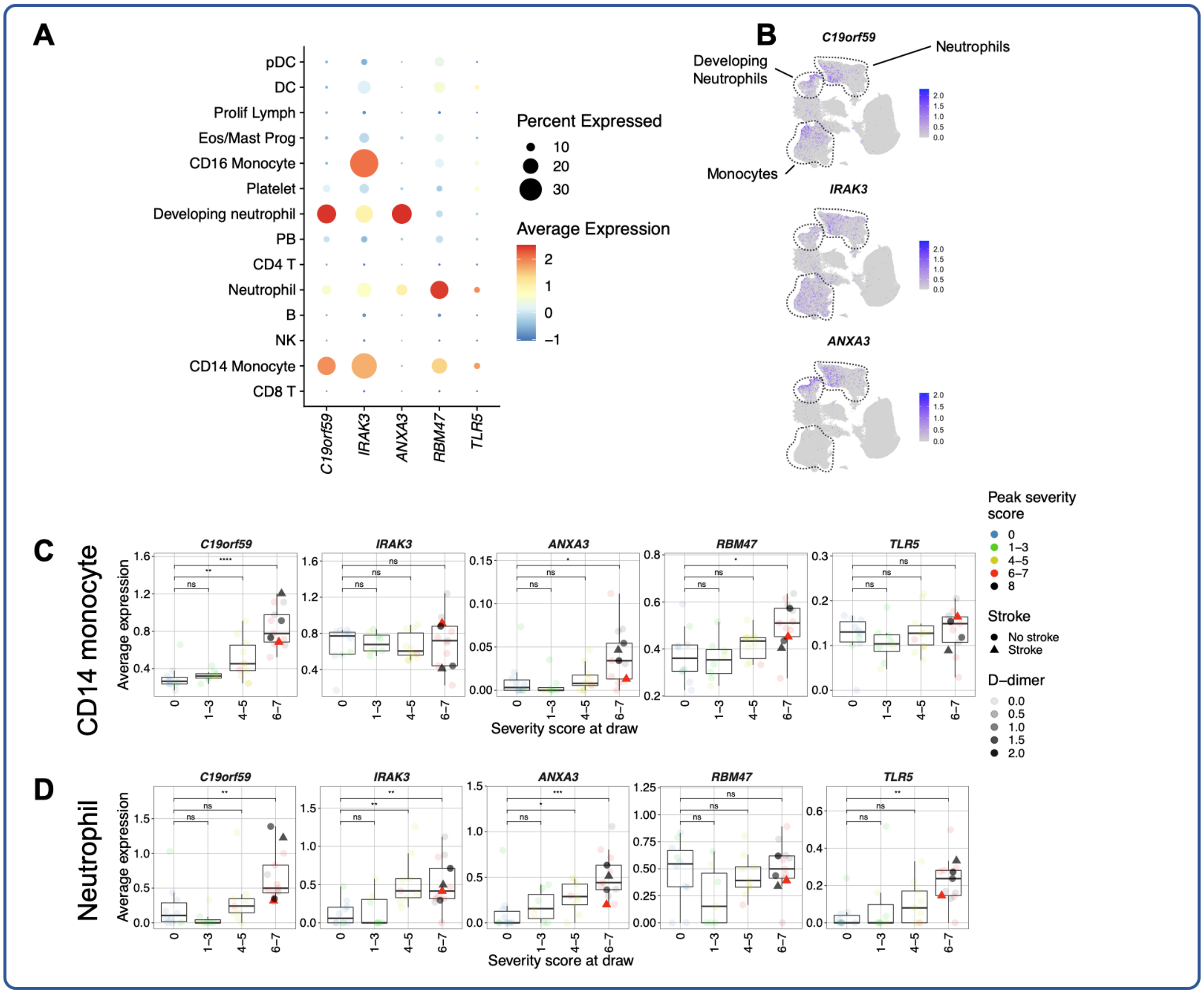


**Fig. S8. Transcripts known to predict stroke risk and outcome are upregulated by monocytes and neutrophils in a disease severity-associated fashion. A**) Dot plot depicting percent and average expression of the top five most predictive genes for stroke risk and outcome reported by Raman, et al. *(47)* by each manually-annotated cell type. **B)** UMAP projection of complete scRNA-seq dataset colored by expression of three stroke-predictive genes that are expressed by developing neutrophils. **C-D)** Box plots depicting average expression of the five stroke-predictive genes from (**A**) in CD14$^{+}$ monocytes (**C**) or canonical neutrophils (**D**). Points are colored by the peak disease severity score and grouped by the disease severity score at time of sample collection. Additionally, point transparency corresponds to measured d-dimer level (μg/mL; known for 5 samples, all other samples are coded as zero), and shape corresponds to a patient experiencing stroke or other thrombotic complication during hospitalization. *, $p < 0.05$; **, $p < 0.01$; ***, $p < 0.001$; ****, $p < 0.0001$, n.s., not significant at $p = 0.05$ by two-sided Wilcoxon rank-sum test with Bonferroni's correction for multiple hypothesis testing.

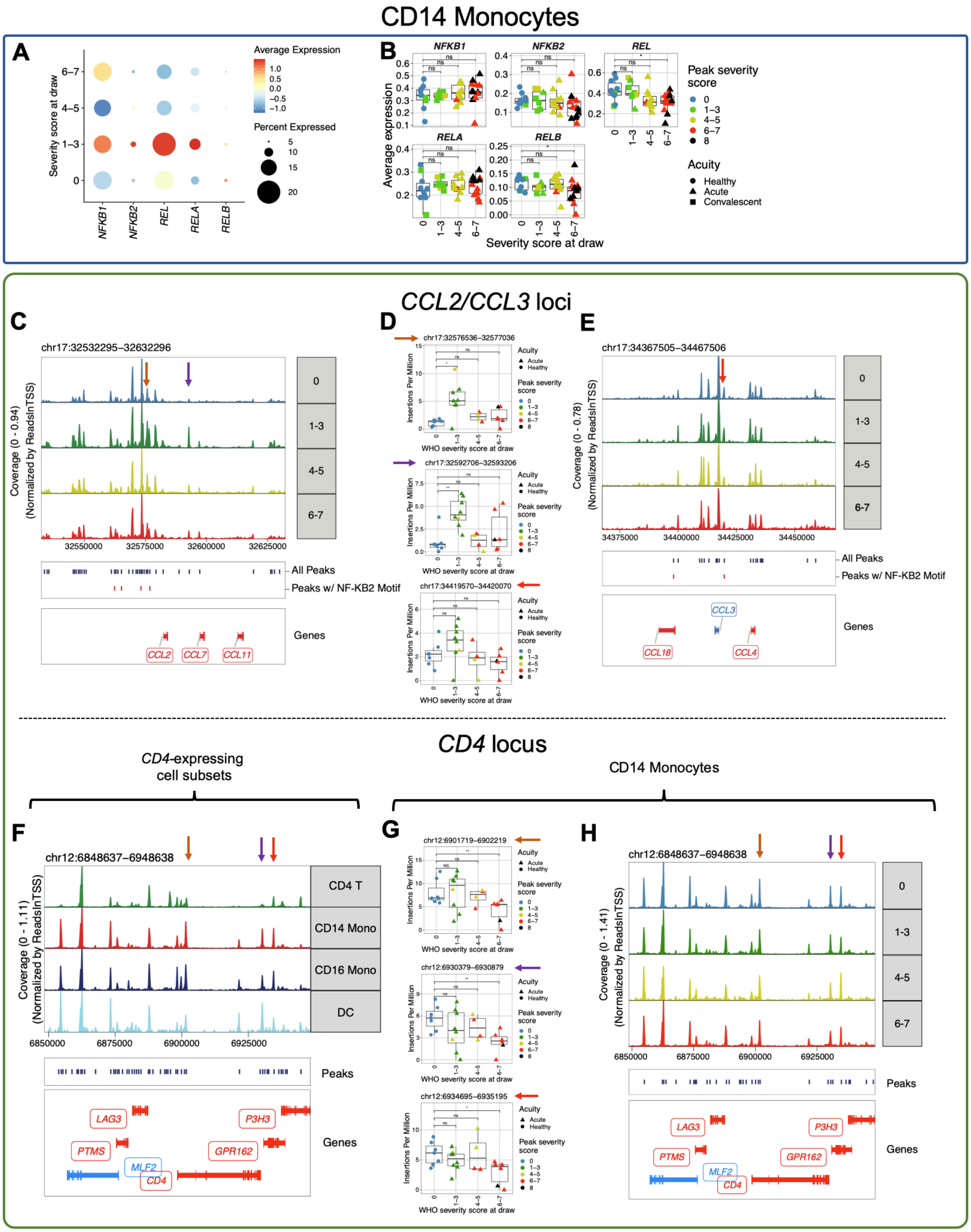


**Fig. S9. Additional analyses of epigenetic regulation in CD14 monocytes in COVID-19. A-B)** Expression patterns of five NF-κB family transcription factors in CD14 monocytes depicted by dot plot showing average and percent expression (**A**) and box plot showing average expression in each COVID-19 sample (**B**). **C,E)** The genome tracks show genomic regions near *CCL2* (**C**) and *CCL3* (**E**) genes. The top panel indicates coverage at different peak regions for CD14 monocytes in different severity groups; the box below shows peaks called from all CD14 monocytes in that region (dark blue) and peaks containing putative strong NF-κB2 binding sites (red); the bottom "Genes" box shows location of *CCL2* (**C**) or *CCL3* (**E**) together with other adjacent genes; blue color means the gene is located on the minus strand, and red color means the gene is located on the plus strand. The arrows indicate peaks of interest whose accessibility is quantified in the corresponding box plots (**D**). **F, H**) The genome tracks show genomic regions near *CD4* gene. The top panel indicates coverage at different peak regions for different cell subsets (**F**) and for CD14 monocytes in different severity groups (**H**); the box below shows peaks called from all PBMCs (**F**) or from the CD14 monocytes (**H**) in that region (dark blue); the bottom "Genes" box shows location of *CD4* and other adjacent genes; blue color means the gene is located on the minus strand, and red color means the gene is located on the plus strand. The arrows indicate monocyte specific peaks with higher accessibility in monocytes and dendritic cells than CD4 T cells. **D,G**) Box plots depicting the Tn5 insertions per million at the peaks marked with the corresponding arrows in CD14 monocytes. Exact p-values for (**D**) top p=0.014 healthy vs. mild, middle p=0.0037 healthy vs. mild. Exact p-values for (**G**) top p=0.0047 healthy vs. severe, middle p=0.0047 healthy vs. severe, bottom p=0.022 healthy vs. mild. Points are colored by the peak disease severity score, shaped according to disease acuity, and grouped by the disease severity score at time of sample collection. *, $p < 0.05$; **, $p < 0.01$; ***, $p < 0.001$; ****, $p < 0.0001$, n.s., not significant at $p = 0.05$ by two-sided Wilcoxon rank-sum test with Bonferroni's correction for multiple hypothesis testing.

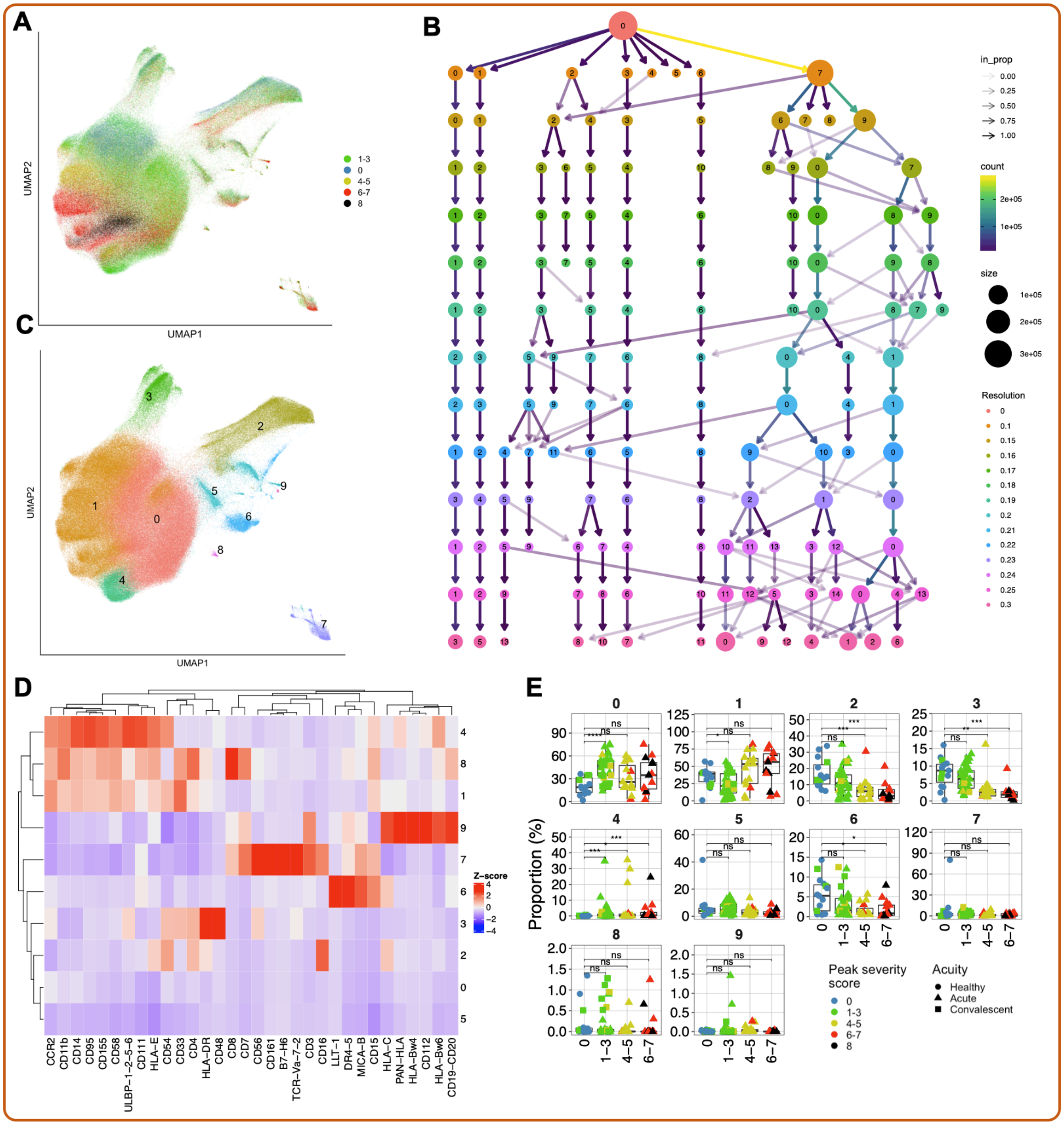


**Fig. S10. CyTOF data reveals reconfiguration of peripheral monocytes in severe COVID-19. A)** UMAP embedding of all peripheral monocytes in our CyTOF dataset. **B)** Cluster tree diagram illustrating the different clusters of monocytes that resulted from performing PARC at different resolution values. A resolution of 0.21 was chosen for unsupervised analysis based on the relative cluster stability achieved at this point. **C)** UMAP embedding of all peripheral monocytes in the PBMC CyTOF dataset colored by PARC cluster. **D)** Heatmap showing the average, z-score normalized protein-level expression of each marker in the PBMC CyTOF panel across each monocyte cluster. **E)** Frequency of each monocyte cluster as a proportion of all monocytes. Points are colored by the peak disease severity score, shaped according to disease acuity, and grouped by the disease severity score at time of sample collection. *, $p < 0.05$; **, p

$< 0.01$; ***, $p < 0.001$; ****, $p < 0.0001$, n.s., not significant at $p = 0.05$ by two-sided Wilcoxon rank-sum test with Bonferroni's correction for multiple hypothesis testing.

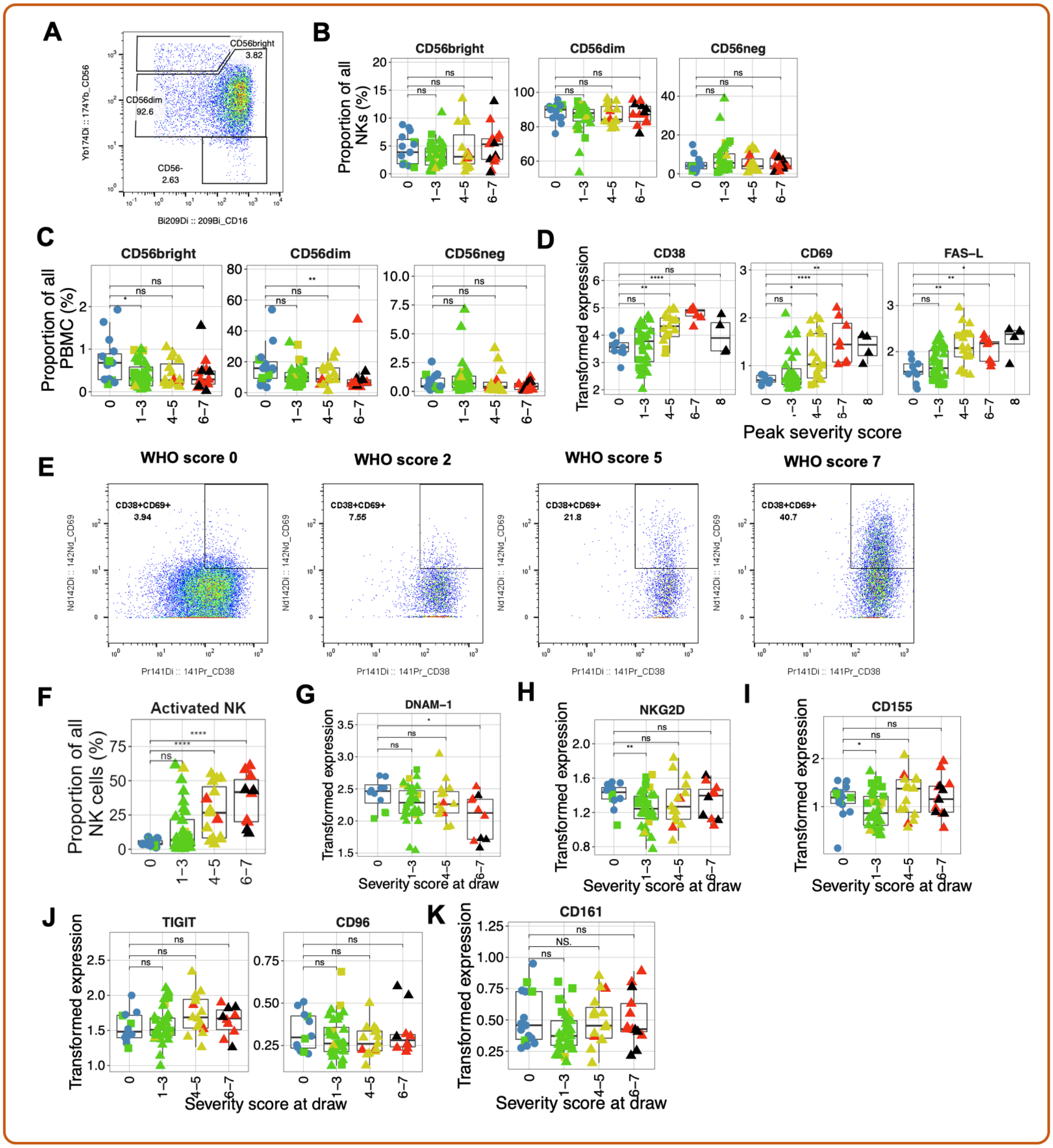


**Fig. S11. Analysis of NK cell subset proportions, markers of activation, and receptors. A)** Representative flow plot showing the gating scheme used to assess the frequencies of NK cell subsets. **B-C)** Boxplots showing the frequency of $CD56^{bright}$, $CD56^{dim}$, and $CD56^{-}$ NK cells as a proportion of NK cells **(B)** and of all PBMC **(C)** in our CyTOF dataset. **D)** Boxplots showing arcsinh-transformed expression of the activation markers CD38, CD69, and Fas-L. **E)** Representative flow plots showing the gating scheme used to identify activated ($CD38^{+}CD69^{+}$) NK cells in patients from each severity bin. **(F)** Boxplot showing the proportion of CD38+CD69+ NK cells in each severity bin. **G-H)** Boxplots showing arcsinh-transformed expression of the activating receptors DNAM-1 (**G**) and NKG2D (**H**) in all NK cells in our

CyTOF dataset. **I)** Boxplot showing arcsinh-transformed expression of DNAM-1 ligand CD155 in the peripheral monocytes of our CyTOF dataset. **J-K)** Boxplots showing arcsinh-transformed expression of the inhibitory receptors TIGIT and CD96/TACTILE (**J**) and CD161 (**K**) in all NK cells in our CyTOF dataset. For all box plots except **(D)**: points are colored by the peak disease severity score, shaped according to disease acuity, and grouped by the disease severity score at time of sample collection. *, $p < 0.05$; **, $p < 0.01$; ***, $p < 0.001$; ****, $p < 0.0001$, n.s., not significant at $p = 0.05$ by two-sided Wilcoxon rank-sum test with Bonferroni's correction for multiple hypothesis testing.

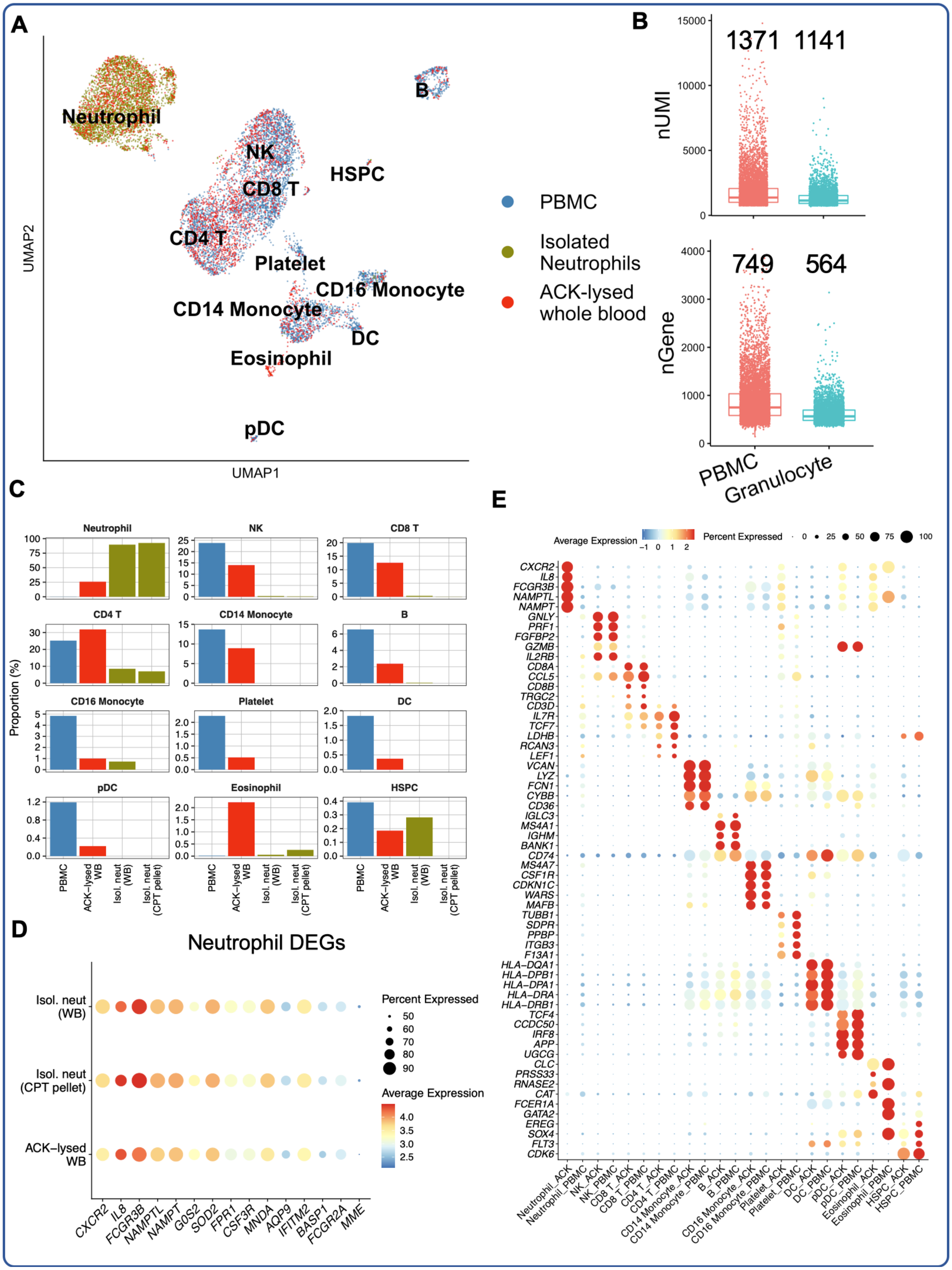

A
UMAP2
UMAP1
Neutrophil
NK
CD8 T
HSPC
B
CD4 T
Platelet
CD16 Monocyte
CD14 Monocyte
DC
Eosinophil
pDC
PBMC
Isolated Neutrophils
ACK-lysed whole blood
B
nUMI
1371 1141
nGene
749 564
PBMC
Granulocyte
C
Proportion (%)
Neutrophil
NK
CD8 T
CD4 T
CD14 Monocyte
B
CD16 Monocyte
Platelet
DC
pDC
Eosinophil
HSPC
PBMC
ACK–lysed WB
Isol. neut (WB)
Isol. neut (CPT pellet)
D
Neutrophil DEGs
Isol. neut (WB)
Isol. neut (CPT pellet)
ACK–lysed WB
Percent Expressed
Average Expression
CXCR2 IL8 FCGR3B NAMPTL NAMPT G0S2 SOD2 FPR1 CSF3R MNDA AQP9 IFITM2 BASP1 FCGR2A MME
E
Average Expression
Percent Expressed

**Fig. S12. Seq-Well enables high-quality single-cell transcriptomic analysis of primary human neutrophils**. Whole blood from a healthy donor was collected into CPT vacutainers, from which PBMCs were isolated and neutrophils were isolated from the PBMC-depleted cell pellet. Additionally, aliquots of whole blood were subjected to neutrophil isolation or red blood cell lysis with ACK buffer. These three cell populations were then analyzed by Seq-Well (see **Methods**). **A)** UMAP projection colored by cell type preparation method. **B)** Box plots showing comparisons of the number of UMIs sequenced (top) and number of genes detected (bottom) in cells annotated to be PBMCs, or cells annotated as granulocytes (neutrophils and eosinophils). The median number of UMIs or genes in each group is plotted above the respective box. The difference in recovered UMIs and gene capture between PBMCs and granulocytes is comparable to that expected by RNA content *(69, 70)*. **C)** Bar plot depicting the proportions of cells from each cell sample preparation method for each annotated cell type. **D)** Dot plot depicting percent and unscaled average expression of the 15 top neutrophil-defining DEGs (see **Table S19**) between the three cell sample preparation methods that yielded neutrophils. **E)** Dot plot depicting average and percent expression of the top five DEGs for each cell type (see **Table S19**) demonstrating comparable expression patterns between PBMCs isolated through centrifugation and PBMC subsets present in ACK-lysed whole blood.

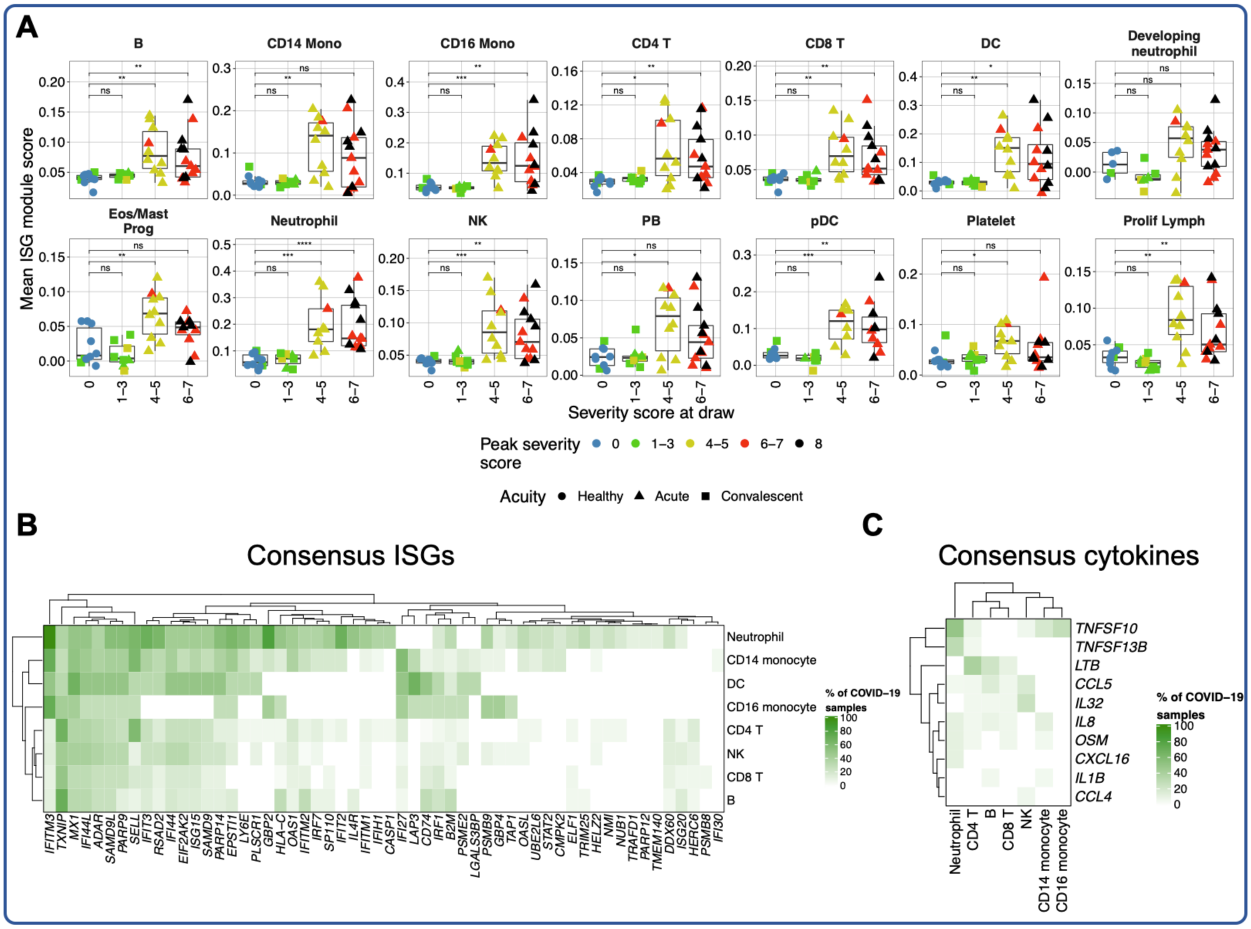


**Fig. S13. Conserved ISG signatures and cytokine-encoding genes upregulated in COVID-19**. **A)** Box plots depicting average expression of ISG module score in each sample from the scRNA-seq dataset in each manually-annotated cell type (see **Methods**, **Table S12**). Points are colored by the peak disease severity score, shaped according to disease acuity, and grouped by the disease severity score at time of sample collection. Boxplot features: minimum whisker, 25th percentile − 1.5 × inter-quartile range (IQR) or the lowest value within; minimum box, 25th percentile; center, median; maximum box, 75th percentile; maximum whisker, 75th percentile + 1.5 × IQR or greatest value within. *, $p < 0.05$; **, $p < 0.01$; ***, $p < 0.001$; ****, $p < 0.0001$, n.s., not significant at $p = 0.05$ by two-sided Wilcoxon rank-sum test with Bonferroni's correction for multiple hypothesis testing. **b,c)** Differential gene expression testing was conducted on 8 major cell types from the scRNA-seq dataset, comparing each COVID-19 sample to the cells of all healthy controls (see **Methods**). The plotted heatmaps depict the percentage of COVID-19 samples in which a given ISG (**B**) or cytokine-encoding gene (**C**) is upregulated in a given cell type.

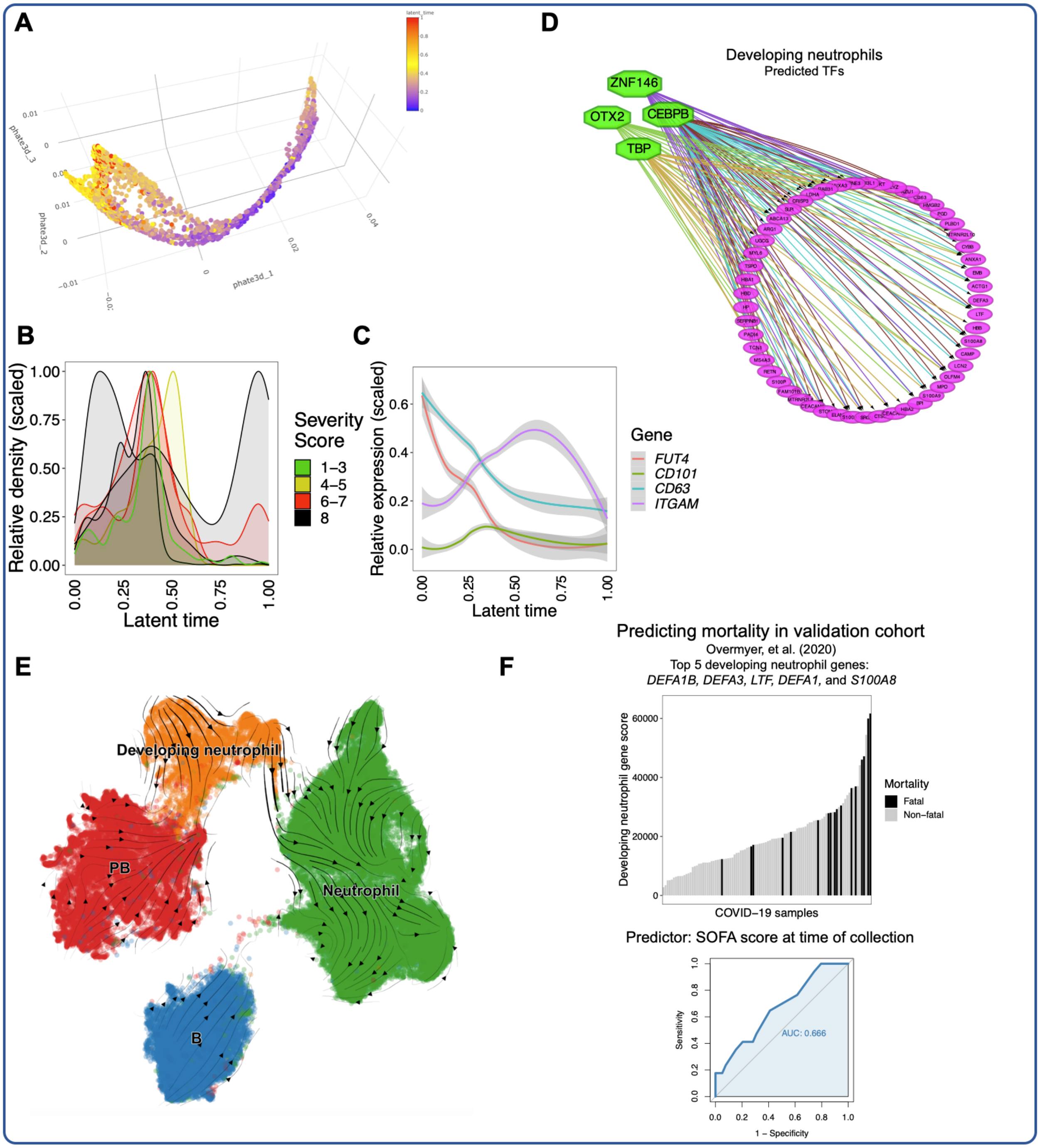


**Fig. S14. Additional analysis of emergency granulopoiesis in severe and fatal COVID-19.** **A)** Three-dimensional PHATE projection of developing neutrophils (see **Supplementary Material**) **B)** Scaled abundances of developing neutrophils present in individual COVID-19 patients across latent time. **C)** Scaled expression of genes reported to define different stages of immature neutrophil development in COVID-19 *(21)*. **D)** Transcription factor activity prediction analysis by iRegulon *(76)*, using positive DEGs for developing neutrophils relative to all other cells as input (**Table S14**). **E)** UMAP projection of developing neutrophils, canonical neutrophils, B cells, and plasmablasts overlaid with RNA velocity stream *(82)*. **F)** Bar plot representing the ranked developing neutrophil signature score (aggregated expression of

*DEFA1B, DEFA3, LTF, DEFA1,* and *S100A8*, see **Methods**) for each COVID-19 samples in bulk transcriptomic dataset published by Overmyer, et al. *(91)*, colored by the 28-day mortality. The corresponding ROC curve is plotted in **Fig. 6**.

**A** **PBMC panel**

| Metal Isotope | Marker | Clone | Manufacturer | Included in pre-mixed panel? |
|---|---|---|---|---|
| 115In | CD3 | UCHT1 | Biolegend | Yes |
| 141Pr | CD161 | DX12 | BD Biosciences | No |
| 142Nd | CD20 | 2H7 | Biolegend | No |
| 142Nd | CD19 | HIB19 | Biolegend | No |
| 143Nd | PAN HLA | W6/32 | Biolegend | Yes |
| 144Nd | CD7 | CD7-6B7 | Biolegend | Yes |
| 145Nd | CD8 | SK1 | Biolegend | Yes |
| 146Nd | CD48 | BJ40 | Biolegend | Yes |
| 148Nd | ICAM1 (CD54) | HA58 | Biolegend | Yes |
| 149Sm | LLT-1 | 402659 | R&D | Yes |
| 151Eu | CD4 | OKT4 | Biolegend | Yes |
| 152Sm | CD15 | W6D3 | Biolegend | No |
| 153Eu | HLA-C | DT9 | Millipore | Yes |
| 154Sm | CCR2 | K036C2 | Biolegend | Yes |
| 155Gd | HLA-E | 3D12 | Biolegend | Yes |
| 156Gd | CD95 | DX2 | Biolegend | Yes |
| 157Gd | Nectin-1 (CD111) | R1.302 | Biolegend | Yes |
| 158Gd | MICA | 159227 | R&D | Yes |
| 158Gd | MICB | 236511 | R&D | Yes |
| 159Tb | DR4 (CD261) | DJR1 | Biolegend | Yes |
| 159Tb | DR5 (CD262) | DJR2-2 | Biolegend | Yes |
| 161Dy | ULBP-1 | 170818 | R&D | Yes |
| 161Dy | ULBP-2 (5 and 6) | 165903 | R&D | Yes |
| 162Dy | TCR Va7.2 | 3C10 | Biolegend | No |
| 164Dy | Nectin-2 (CD112) | TX31 | Biolegend | Yes |
| 165Ho | CD155 (PVR) | SKII.4 | Biolegend | Yes |
| 166Er | HLA-Bw4 | REA274 | Miltenyi | Yes |
| 168Er | HLA-Bw6 | REA143/ 130-095-212 | Miltenyi | Yes |
| 169Tm | CD14 | M5E2 | Biolegend | Yes |
| 170Er | CD11b | ICRF44 | Biolegend | Yes |
| 171Yb | LFA-3 (CD58) | TS2/9 | Biolegend | Yes |
| 172Yb | CD33 | WM53 | Biolegend | Yes |
| 174Yb | CD56 | NCAM16.2 | BD Pharmingen | Yes |
| 176Yb | B7-H6 | 875001 | R&D | Yes |
| 209Bi | CD16 | CD16 | Fluidigm | No |
| 89Y | HLA-DR | L243 | Biolegend | Yes |

**B** **NK panel**

| Metal Isotope | Marker | Manufacturer | Clone | Included in pre-mixed panel? |
|---|---|---|---|---|
| 89Y | CD57 | Biolegend | HCD57 | Yes, surface panel |
| Qdot | HLA-DR | Life technologies | Tu36 | No |
| 115In | CD3 | Biolegend | UCHT | Yes, surface panel |
| 141Pr | CD38 | Biolegend | HIT2 | Yes, surface panel |
| 142Nd | CD69 | Biolegend | FN50 | Yes, surface panel |
| 143Nd | CD33 | Biolegend | WM53 | Yes, surface panel |
| 143Nd | CD14 | Biolegend | M5E5 | Yes, surface panel |
| 144Nd | CD2 | Biolegend | RPA-2.10 | Yes, surface panel |
| 145Nd | LILRB1 | Biolegend | GHI/75 | No |
| 146Nd | CD19 | Biolegend | HIB19 | Yes, surface panel |
| 147Sm | CD8 | Biolegend | SK1 | Yes, surface panel |
| 149Sm | CD4 | Biolegend | SK3 | Yes, surface panel |
| 151Eu | CD62L | Biolegend | DREG-56 | Yes, surface panel |
| 153Eu | KIR2DS4 | R&D systems | 179315 | Yes, surface panel |
| 154Sm | KIR2DS2 | Abcam | Poly-clonal | Yes, surface panel |
| 155Gd | NKp46 | Biolegend | 9.00E+02 | Yes, surface panel |
| 156Gd | NKG2D | Biolegend | 1D11 | Yes, surface panel |
| 157Gd | TIGIT | R&D systems | 741182 | Yes, surface panel |
| 158Gd | 2B4 | Biolegend | C1.7 | Yes, surface panel |
| 159Tb | DNAM-1 | BD biosciences | DX11 | Yes, surface panel |
| 160Gd | FAS-L | Biolegend | NOK-1 | Yes, surface panel |
| 161Dy | NKp30 | Biolegend | P30-15 | Yes, surface panel |
| 162Dy | Siglec-7 | Biolegend | S7.7 | Yes, surface panel |
| 163Dy | NKG2C | R&D systems | 134522 | Yes, surface panel |
| 164Dy | NKp44 | Biolegend | P44-8 | Yes, surface panel |
| 165Ho | TACTILE | Biolegend | NK92.39 | Yes, surface panel |
| 166Er | KIR2DL1/KIR2DS5 | R&D systems | 143211 | Yes, surface panel |
| 167Er | CD94 | Biolegend | DX22 | Yes, surface panel |
| 168Er | CXCR6 | Biolegend | K041E5 | Yes, surface panel |
| 169Tm | PD1 | Biolegend | EH12.2H7 | Yes, surface panel |
| 170Er | KIR2DL5 | Miltenyi | UP-R1 | Yes, surface panel |
| 171Yb | NKG2A | R&D systems | 131411 | Yes, surface panel |
| 172Tb | NTB-A | Biolegend | NT-7 | Yes, surface panel |
| 173Yb | KIR3DL1 | BD biosciences | DX-9 | Yes, surface panel |
| 174Yb | CD56 | BD biosciences | NCAM16.2 | Yes, surface panel |
| 175Lu | KIR2DL3 | R&D systems | 180701 | Yes, surface panel |
| 209Bi | CD16 | Fluidigm | 3G8 | No |
| 148Nd | FcRg | Millipore | polyclonal | Yes, IC panel |
| 150Nd | Syk | Biolegend | 4D10.2 | Yes, IC panel |
| 152Sm | Ki-67 | Biolegend | Ki-67 | Yes, IC panel |
| 176Yb | Perforin | Abcam | B-D48 | Yes, IC panel |

**C** **Whole PBMC**

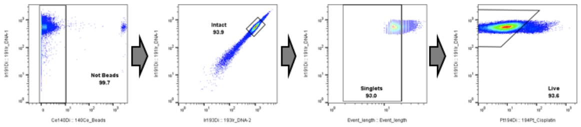

**D** **Purified NK cells**

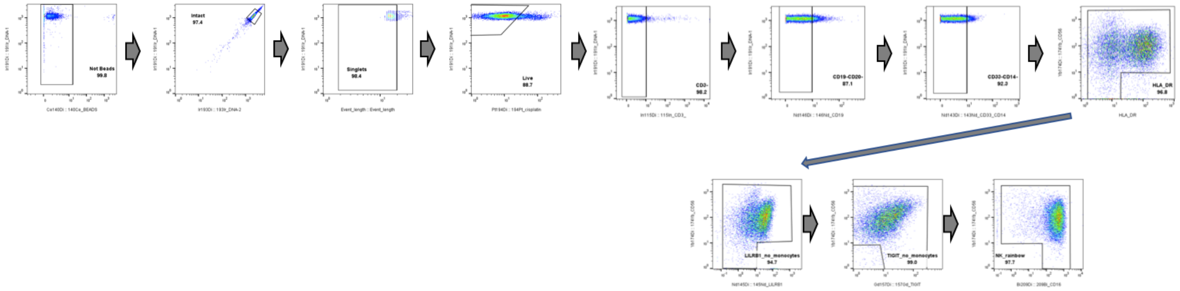

**E**

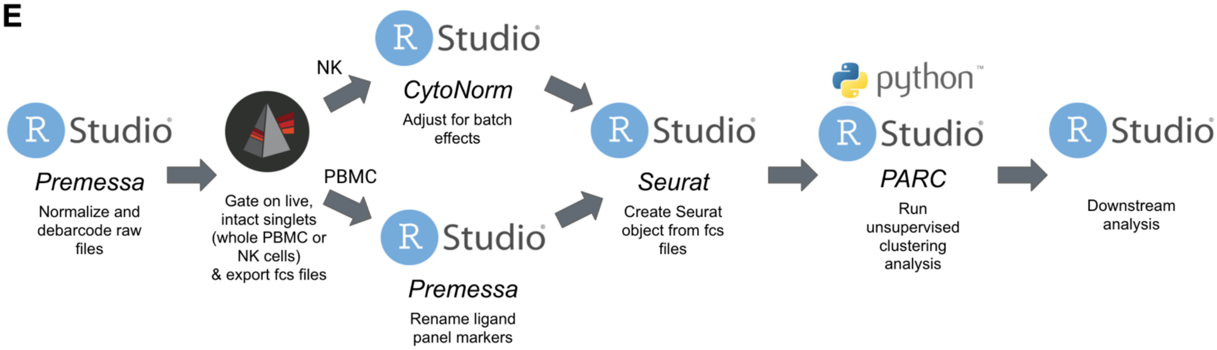

**Fig. S15. CyTOF panels and preprocessing workflow**. **A-B)** Tables showing the antibodies included in the PBMC (**A**) and NK (**B**) CyTOF panels **(**see **Methods)**. **c-d)** Gating strategies used to identify live, intact singlets (**C**) and live, intact, singlet NK cells (**D**). **E)** Diagram outlining the preprocessing steps utilized for each CyTOF dataset prior to analysis.
